# A replicated patient-specific component of tumour telomere length across two pan-cancer cohorts

**DOI:** 10.64898/2026.08.31.748414

**Authors:** Subhajit Dutta

## Abstract

Bulk telomere length measured from tumour sequencing is routinely read as a property of the cancer cells. A tumour specimen is a mixture, however, and the patient who supplies it has a telomere length of their own. Here I re-analyse published pan-cancer telomere estimates and ask how much of a tumour’s telomere length is the patient’s.

A calibration step comes first. Whole-genome and low-pass estimates recover the known cross-sectional attrition of leukocyte telomeres with age (−26.6 bp per year in blood normals), whereas whole-exome estimates do not: adjusted for cancer type, sequencing centre and sex the exome slope is −0.6 bp per year, and in 684 blood-normal aliquots sequenced by both assays the whole-genome estimate declines at −38.9 bp per year while the exome estimate from the same DNA does not decline at all (difference −41.5 bp per year, *P* = 3 × 10⁻¹⁰). Because exome data are 78.6% of the resource, downstream analyses use only the calibrated libraries.

Within those, tumour telomere length tracks the patient’s matched normal (Spearman ρ = +0.395; 22 of 23 cancer types positive), and this replicates in a second consortium measured with a different estimator (PCAWG ρ = +0.472, 24 of 24 histologies). Entering cancer type, sequencing centre and library type as fixed effects leaves β = 0.385, and the coefficient is unmoved by age, sex, purity, leukocyte fraction, ploidy, coverage and continental ancestry (range 0.406–0.429). Pure normal-cell admixture is rejected as the sole explanation: the mixture model requires β_H = 1 and β_H×p = −1, and both are rejected jointly (*P* = 0.001).

Tumour purity, leukocyte fraction and age each explain only ∼1–3% of within-cohort variance, and none alters the cross-cancer ranking. The between-cohort coefficient is not interpretable: it appears to show one-to-one correspondence with tissue-of-origin telomere length, but that value depends on which tissue supplies the reference and on the spread of the predictor, and falls to 0.44 with an organ-matched solid-tissue normal.

Bulk tumour telomere length is therefore a composite phenotype containing a replicated patient-specific component. Telomere biomarker studies should carry matched normal telomere length as a covariate rather than treating tumour telomere length as tumour-intrinsic.

## Introduction

Telomeres are nucleoprotein caps that shorten with each somatic division, because the replication machinery cannot copy a linear chromosome to its end [1], and are protected from the DNA-damage machinery by the shelterin complex [2]. Their progressive erosion is one of the better-characterised molecular correlates of ageing in human tissues [3,4], and telomere dysfunction sits at the junction between replicative senescence, genome instability and malignant transformation [5]. Sustained proliferation therefore requires a telomere maintenance mechanism, and its acquisition is a recognised enabling characteristic of cancer [6].

Two such mechanisms dominate. Most tumours reactivate telomerase [7], frequently through recurrent promoter mutations that create de novo transcription-factor binding sites at *TERT* [8,9], which are especially common in glioma and other tumours arising from slowly self-renewing tissue [10]. A minority instead use alternative lengthening of telomeres (ALT), a recombination-based route [11] strongly associated with inactivation of the chromatin remodeller *ATRX* or its partner *DAXX* [12,13]. Telomere length itself feeds back on this machinery: telomere-length-dependent looping by non-telomeric TRF2 regulates *TERT* transcription [14], and the same shelterin protein transduces telomere length into altered expression of immune receptors within the tumour microenvironment [15]. Telomere state is thus not merely an outcome of proliferation but an input to the regulatory circuitry of the cell.

Against this background, two consortium-scale surveys established what is now the reference description of telomere length across human cancer. Barthel and colleagues estimated telomere length in 18,430 TCGA samples spanning 31 cancer types and reported two headline observations: telomeres are shorter in tumours than in matched non-neoplastic tissue — generalising an observation with a long single-cohort history, reviewed in [16] — and they are longest in sarcoma and glioma [17]. Sieverling and colleagues, analysing whole genomes from the Pan-Cancer Analysis of Whole Genomes consortium [18], described the genomic footprints that distinguish telomerase-driven from ALT-driven tumours [19]. Both rest on computational estimation of telomere content from short-read sequencing — an approach implemented in several tools [20,21,22,23] whose behaviour across library preparations has received less scrutiny than the biology built upon it.

There is a second body of literature that these surveys do not usually meet. Leukocyte telomere length is among the most studied constitutional traits in human epidemiology. It shortens with age at a rate that has been meta-analysed across hundreds of thousands of individuals [24,25], it is substantially heritable [26], its common-variant architecture has been mapped in biobank-scale genome-wide association studies [27], and Mendelian randomisation using those variants implicates longer genetically-predicted telomeres in the risk of several cancers [28]. Telomere length also varies systematically between tissues within the same individual while remaining correlated across them [29], and it is a recognised axis of organismal ageing biology [30,31].

These two literatures make an assumption that has not been tested directly. Pan-cancer telomere surveys treat a tumour’s telomere length as a property of the tumour, to be explained by its mutations and its maintenance mechanism. But every tumour specimen is a mixture of malignant cells, stroma and leukocytes, and it is supplied by a person who has a constitutional telomere length, a tissue of origin with its own characteristic telomere length, and an age. Tumour purity varies enormously across TCGA cohorts and is known to confound molecular inference broadly [32,33], as does immune and stromal infiltration [34,35]. Whether any of this reaches the telomere landscape is unknown, and the one report that examined it — in prostate cancer — noted in a single sentence that tumour telomere lengths appeared independent of purity [36].

I set out to test whether the pan-cancer telomere landscape is confounded by specimen composition, and found that it is not. What emerged instead was a different and more tractable effect: a measurable, replicated component of a tumour’s telomere length is the patient’s own. This paper reports that component, the calibration work required to make it trustworthy, the boundary of what the data can and cannot identify about its tissue-level counterpart, and the several hypotheses that did not survive testing.

## Results

### The analysis set

All tumour–normal and composition analyses below use whole-genome (WGS) and low-pass whole-genome (LPS) telomere estimates; the exome data are analysed only to establish that restriction. Three sample counts recur and are distinguished here once. The published paired set contributes **1,942** tumour/normal pairs on these libraries, all with a telomere estimate for both members; this is what the joint distributions in Fig. 1a, 1b show. Restricting to cancer types with at least 25 such pairs drops 7 pairs and leaves **1,935** in 23 types, which is the paired analysis set used for every tumour-versus-normal model. The cross-cancer composition analyses additionally require a copy-number purity call and are therefore run on primary tumours only: 1,791 primary tumours on these libraries, 1,728 with a purity call, and **1,706** in the 21 cancer types with at least 25 such tumours. That set defines the cross-cancer landscape analyses. Paired sensitivity analyses that combine a matched normal with a composition covariate are a third set again, since they require both, and their sizes are stated where they occur. The paired set exceeds the primary-tumour set because it also contains metastatic and blood-derived tumours. The exclusion of whole-exome (WXS) estimates is not a preference but a consequence of the calibration result presented in the next section (Fig. 2). The full sample-selection cascade is given in Supplementary Fig. S1: of 18,430 samples with a telomere estimate, 3,947 were generated by WGS or LPS, of which 1,791 are primary tumours, 1,728 match a copy-number-derived purity call, and 1,706 fall in cancer types with at least 25 such samples (21 of 23 types; SKCM and DLBC were excluded at n = 15 and n = 7). No join lost more than 4% of tumours.

**Figure 1.**
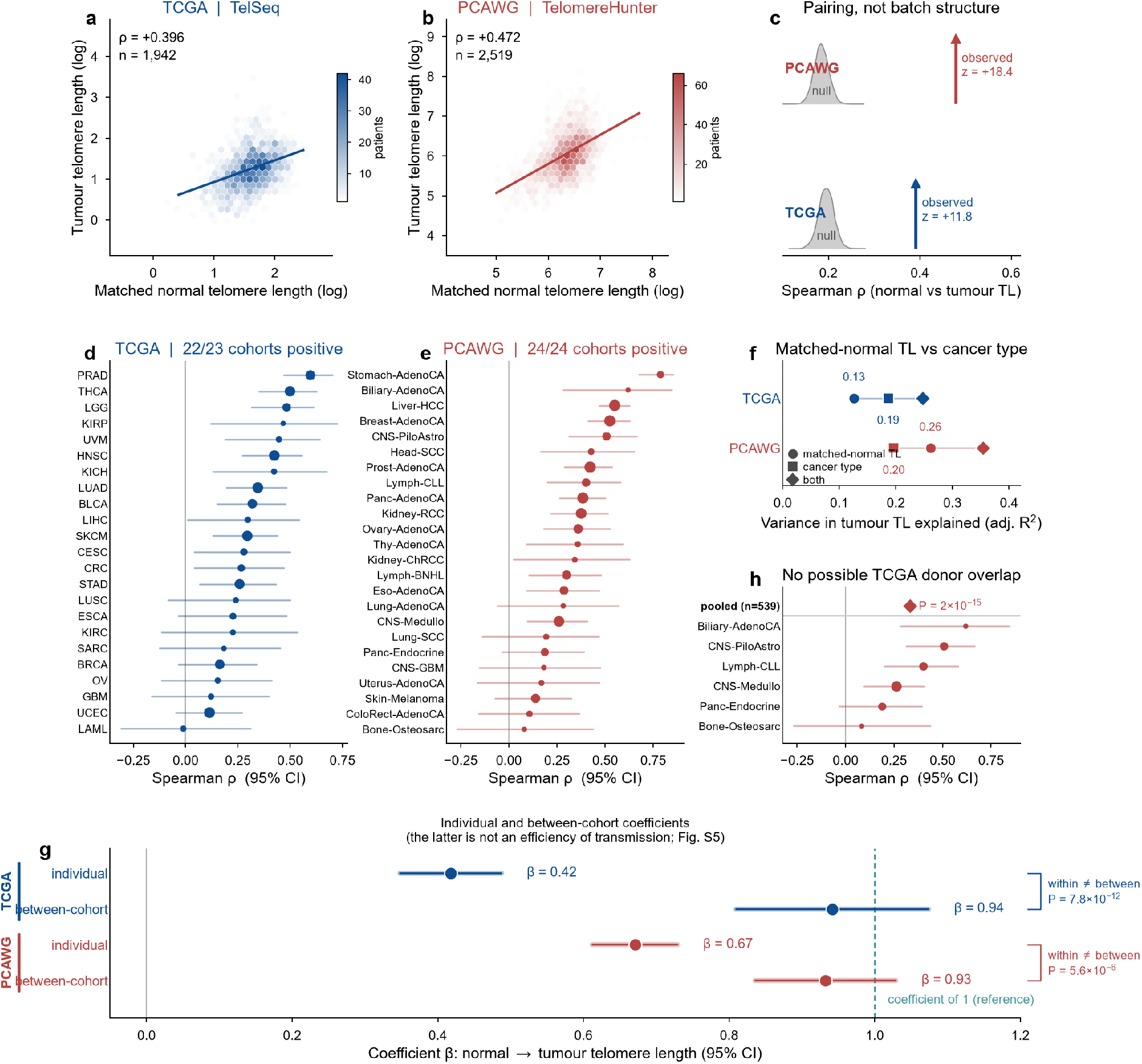
Tumour telomere length carries a replicated component of the patient’s own telomere length. (**a**, **b**) Joint density of matched-normal and tumour telomere length in TCGA (TelSeq, n = 1,942 — the complete paired whole-genome and low-pass set, before the minimum-cohort-size rule that defines the 1,935-pair analysis set used in the models) and PCAWG (TelomereHunter, n = 2,519); lines are least-squares fits over the central 99% of the data. (**c**) Permutation test: normals were shuffled among patients within cancer type × sequencing centre × library type strata (TCGA) or histology strata (PCAWG), 10,000 permutations. Grey densities are the nulls; arrows mark the observed values. (**d**, **e**) Per-cohort Spearman correlations with percentile bootstrap 95% confidence intervals; point size is proportional to cohort size. (**f**) Variance in tumour telomere length explained by the patient’s own normal telomere length, by cancer type, and by both. (**g**) Within/between decomposition: the individual-level (within-cohort) and between-cohort coefficients with the equality test for each cohort. The dashed line marks a slope of one; as shown in Supplementary Fig. S5, the between-cohort coefficient is not interpretable as an efficiency of transmission and is plotted here for completeness. (**h**) Restriction to 13 PCAWG histologies with no TCGA counterpart, so that donor overlap is impossible.

**Figure 2.**
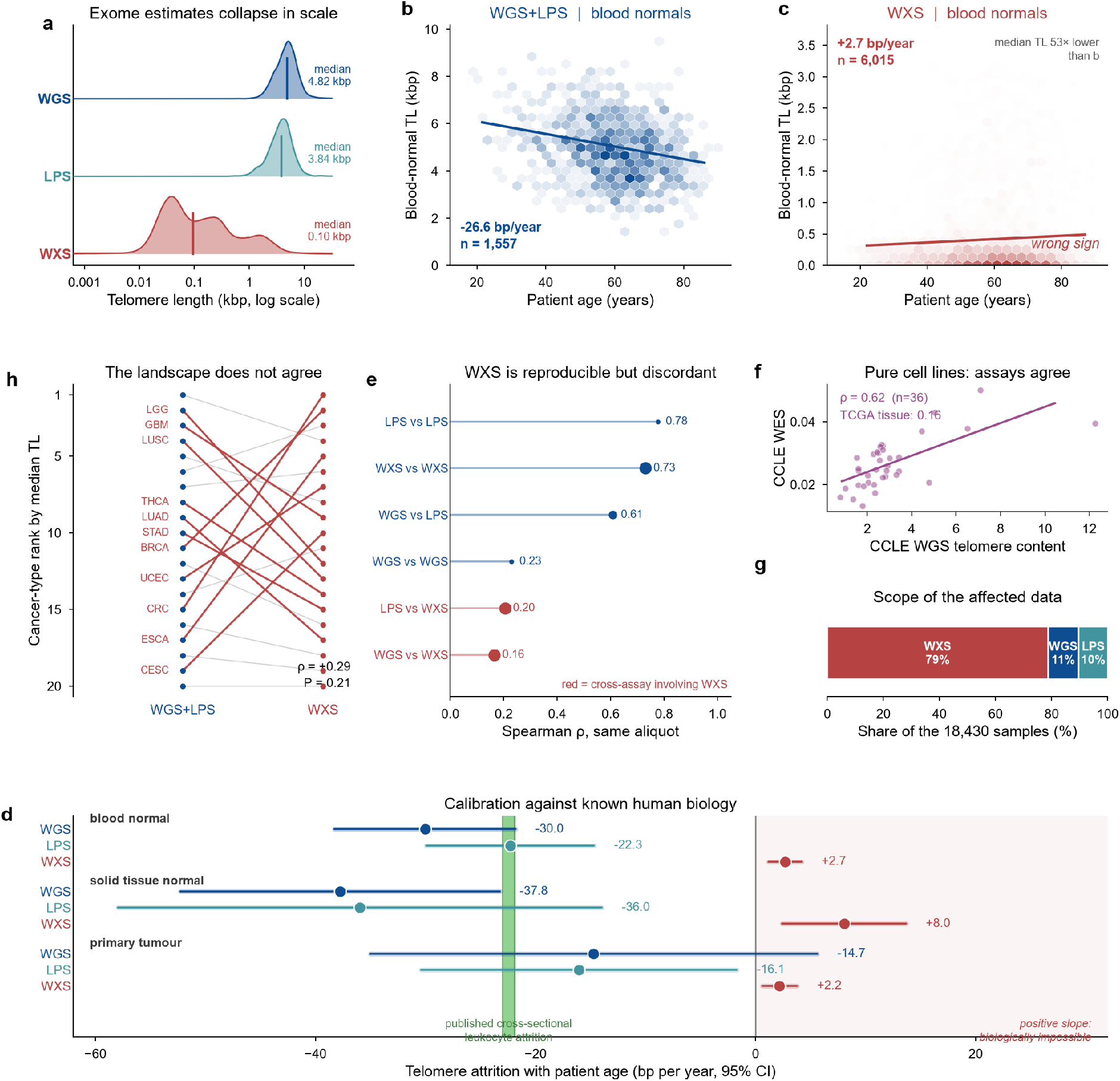
Whole-genome and low-pass telomere estimates are calibrated against age; whole-exome estimates are not. (**a**) Telomere-length distributions by assay, all sample types pooled; medians annotated. (**b**, **c**) Blood-normal telomere length against patient age for WGS+LPS and for WXS, with least-squares fits; note the axis scales differ. (**d**) Telomere attrition with age by sample type and assay, with 95% confidence intervals. The green band spans the two published cross-sectional meta-analytic estimates for leukocyte telomere attrition; the shaded half-plane marks positive slopes. Unadjusted exome slopes fall in it, but the exome estimate is better described as carrying no age signal: adjusted and within-cohort estimates are indistinguishable from zero (Supplementary Table S7b). (**e**) Spearman correlation between telomere estimates from the same physical aliquot, by assay pair. (**f**) CCLE cell lines: WGS versus WES telomere content. (**g**) Assay composition of the 18,430-sample resource. (**h**) Cross-cancer ranking derived from WXS against that derived from WGS+LPS; cohorts shifting five or more places are highlighted.

### Tumour telomere length tracks the patient’s own normal telomere length

Barthel’s paired set provides, for the same patient, a tumour telomere length and a matched non-neoplastic telomere length. These are strongly associated across all 1,935 pairs in the paired analysis set (Spearman ρ = +0.395, *P* = 2.2 × 10⁻⁷³), and the association is positive in 22 of 23 cancer types with a median within-cohort ρ of +0.28 (Fig. 1a, 1d). The magnitude is close to the ρ = 0.37 reported in prostate cancer [36], and the present prostate estimate (ρ = 0.60) is the strongest of any cohort.

Because a tumour and its matched normal are sequenced at the same centre with the same library preparation, batch structure alone could generate this correlation. To test that, normals were permuted among patients *within* strata defined by cancer type, sequencing centre and library type — preserving those three factors while destroying the patient pairing. The test is restricted to the 1,926 pairs that fall in a stratum containing at least eight pairs, on which the observed correlation is ρ = +0.392. Across 10,000 permutations the null distribution centred on ρ = +0.195 (SD 0.017), placing the observation 11.8 standard deviations above the null (*P* < 10⁻⁴; Fig. 1c). Preserved cohort and technical structure therefore generates a positive null on its own, but it cannot account for the matched-patient excess. It does not license an additive split of the observed correlation into batch and pairing components: correlations are not additive, and the null mean is not the batch share. The magnitude that survives removing that structure directly is given below.

The permutation establishes that the observed pairing exceeds what shared technical structure can generate, but it does not give the adjusted magnitude. Entering that structure directly as fixed effects does. Regressing tumour on matched-normal log telomere length with a fixed effect for cancer type gives β = 0.417 (95% CI 0.352–0.483); adding sequencing centre and library type gives 0.407 (0.339– 0.474); and a single fixed effect for the complete cancer-type × centre × library-type stratum — the same unit the permutation shuffled within, 43 strata in all, of which the permutation used the 40 containing at least eight pairs — gives **β = 0.385 (0.316–0.455, *P* = 7.6 × 10⁻²⁷)**, with a partial *R*² of 0.059. Residualising both variables on that complete stratum and correlating the residuals gives Spearman ρ = +0.280 (*P* = 3.1 × 10⁻³⁶). The direct adjustment and the shuffling therefore agree: removing the measured shared technical structure attenuates the association by roughly a quarter and leaves it strongly present (Fig. 5d, Supplementary Table S18).

The finding replicates. In PCAWG, where telomere content was measured with TelomereHunter rather than TelSeq and the donor set is largely non-TCGA, the matched-patient correlation is ρ = +0.472 (*P* = 3.9 × 10⁻¹⁴⁰, n = 2,519), positive in 24 of 24 histologies meeting the same minimum cohort size, with median ρ = +0.34, and 18.4 standard deviations above its own stratified null (computed on the 2,397 specimens in adequately sized histology strata; Fig. 1b, 1e). PCAWG shares approximately 800 donors with TCGA; restricting to 13 histologies that have no TCGA counterpart at all, so that donor overlap is impossible, leaves ρ = +0.332 (*P* = 2.3 × 10⁻¹⁵, n = 539), positive in 6 of 6 cohorts with ≥ 20 specimens (Fig. 1h).

Matched-normal telomere length is not a weak predictor relative to cancer type. This quantity mixes patient-level and cohort-level structure and is not the individual-level component isolated later. On the paired TCGA set, the patient’s own normal telomere length explains 12.6% of the variance in tumour telomere length against 18.6% for cancer type, and the combined model explains 24.7%, exceeding either predictor alone. In PCAWG matched-normal telomere length explains 26.1% against 19.6% for histology (Fig. 1f).

### Whole-exome telomere estimates fail to recover the expected age attrition

Any claim built on these estimates depends on the estimates meaning something. Leukocyte telomere length declines with age at a rate established in two independent meta-analyses, with cross-sectional estimates of −23 bp per year [25] and −21.9 bp per year [24]; TCGA is a cross-sectional cohort, so the cross-sectional figure is the appropriate comparator. Barthel’s blood normals are essentially pure leukocyte DNA at a known patient age and therefore provide a direct calibration.

WGS and LPS pass it. Blood-normal telomere length declines at −30.0 bp per year (WGS, n = 730) and −22.3 bp per year (LPS, n = 827), pooling to −26.6 bp per year (*P* = 1.1 × 10⁻¹⁸) — squarely within the published cross-sectional range (Fig. 2b, 2d). Solid-tissue normals decline more steeply still (−37.8 and −36.0 bp per year). WXS fails it. Blood normals measured by WXS return **+2.7 bp per year** unadjusted (*P* = 6.0 × 10⁻⁴) and solid normals **+8.0 bp per year** (*P* = 5.2 × 10⁻³) (Fig. 2c, 2d). The positive sign is not itself robust, and it is worth saying so: adjusting for cancer type, sequencing centre and sex moves the exome blood slope to −0.6 bp per year (95% CI −2.1 to +0.9, *P* = 0.45), and estimating it inside each cancer type and pooling by inverse-variance meta-analysis gives −0.14 bp per year (*P* = 0.056). The same adjustments leave the calibrated assays essentially unchanged (WGS −29.4, LPS −28.9 bp per year adjusted; −33.1 and −27.3 within-cohort). What the exome estimates fail at is not the sign but the recovery of the age signal, which is the more damaging finding: the single best-established quantitative fact about telomeres in human tissue is absent from them.

The decisive test holds the specimen constant. For 684 blood-normal aliquots the identical DNA was sequenced by both assays, so patient, age, cancer type, centre and material are the same by construction and only the assay differs. On those aliquots the whole-genome estimate declines at −38.9 bp per year (95% CI −47.8 to −30.0, *P* = 5.4 × 10⁻¹⁷) while the exome estimate does not decline at all (+2.6, 95% CI −6.7 to +11.9, *P* = 0.59); the within-aliquot difference in slope is −41.5 bp per year (*P* = 3.3 × 10⁻¹⁰) (Supplementary Table S7b). Cohort composition, centre and sex cannot account for this, because none of them varies between the two measurements being compared. The exome estimates are also on a different scale entirely — a median of 0.10 kbp against 4.82 kbp for WGS, a 53-fold difference in blood normals (Fig. 2a). Same-aliquot replicate pairs confirm the discordance: WXS is internally reproducible (WXS vs WXS ρ = 0.73, n = 1,273) yet agrees poorly with WGS on the same physical aliquot (ρ = 0.165, n = 2,040), while WGS and LPS agree well (ρ = 0.61) (Fig. 2e). Barthel disclosed that exome-based estimates were less robust than genome-based ones; the calibration test quantifies how much less, and shows that what fails is the recovery of the age signal itself.

Two further observations bound the problem. First, it was not reproduced in cell lines: across 36 CCLE lines — pure material with clean DNA — WGS and WES telomere estimates agree reasonably (ρ = 0.62) [37], which suggests the problem is context dependent rather than intrinsic to exome capture (Fig. 2f). Second, it matters: WXS constitutes 78.6% of the 18,430-sample resource (Fig. 2g), and the cross-cancer ranking derived from WXS does not agree with the ranking derived from WGS and LPS (ρ = +0.29, *P* = 0.21; Fig. 2h).

### Pure normal-cell admixture cannot explain the association

The obvious deflationary explanation is that the tumour specimen physically contains the patient’s own normal cells, so their telomeres correlate by construction. This is falsifiable. Under pure admixture, measured telomere length is *p*·TL_tumour + (1 − *p*)·TL_host, so the regression coefficient of measured on host telomere length must equal (1 − *p*) and vanish as purity approaches 1. That identity is a statement about raw telomere length, so the test is run on the raw scale, where the predicted coefficient applies directly; on the log scale used elsewhere the expectation is not (1 − *p*) but (1 − *p*)·TL_host/TL_measured, and is derived rather than assumed.

Stratifying by copy-number-derived purity quartile [38] does not reproduce that behaviour (Fig. 4a). On the raw scale the coefficient is 0.426 in the lowest quartile (median purity 0.38) and 0.360 in the highest (median purity 0.90), a ratio of **0.85**, where pure admixture requires 0.16 — a fall of 15% against a required fall of 84%. The quartile prediction uses median purities and is an approximation, so the formal test is the continuous model. Under Y = *p*T + (1 − *p*)H with tumour and host telomere length independent, that model predicts β_H = 1 and β_H×p = −1, not merely a zero interaction. Observed: β_H = 0.435 (*P* = 9.4 × 10⁻⁴ against 1) and β_H×p = −0.092 (*P* = 2.6 × 10⁻⁴ against −1); the joint Wald test rejects the pure-admixture null outright, F(2, 1820) = 6.87, **\*P* = 0.0011** (n = 1,824). The rejection does not depend on modelling the main effect of purity linearly: with a natural cubic spline in purity it is F = 6.10, *P* = 0.0023. The interaction itself is not significantly different from zero (*P* = 0.71), which is a separate statement. On the log scale used elsewhere in this work the correctly derived prediction is 0.21 and the observed ratio is 1.26. Both scales agree: the coefficient does not decline anything like steeply enough for admixture alone to generate it. This falsifies pure admixture as the sole explanation; it does not exclude some admixture contribution coexisting with a patient-specific association, and no such exclusion is claimed. Across cohorts the same conclusion holds: the size of the host effect does not track a cohort’s median purity (ρ = −0.09, *P* = 0.67; Fig. 4d).

The association is likewise present across telomere-maintenance mechanisms (ATRX/DAXX-truncating ρ = 0.42, TERT-modified ρ = 0.47, other ρ = 0.47; Fig. 4b), across assays (TCGA WGS 0.28, TCGA LPS 0.42, PCAWG WGS 0.47; Fig. 4c) and across tumour stages (ρ = 0.35, 0.46, 0.41 and 0.32 for stages I–IV; Fig. 4g). These are stratified estimates, not formal tests of interaction. It survives nested adjustment for purity, leukocyte fraction, age, ploidy and coverage, falling only from β = 0.434 to β = 0.390 (all *P* < 10⁻³²; Fig. 4f).

### Purity, immune composition and reference tissue do not explain the landscape

Having established what does contribute, it is worth being explicit about what does not, because these are the explanations the field would reach for first.

### Purity

Within cancer types, telomere length declines weakly with purity: 17 of 21 cohorts negative (sign test *P* = 0.0072), median ρ = −0.11, corresponding to about 1% of within-cohort rank variance, with 4 of 21 significant after Benjamini–Hochberg correction [39] (Fig. 3a). The pooled effect is real (β = −0.253, 95% CI −0.370 to −0.135) but it does not propagate: re-ranking cancer types by purity-adjusted marginal means leaves the ranking essentially unchanged (Spearman ρ = 0.96 between raw and adjusted rankings; mean absolute rank shift 1.0 places). Sarcoma moves from rank 1 to rank 3 and lower-grade glioma from 2 to 1 (Fig. 3b). Barthel’s second headline claim is therefore robust to purity adjustment.

**Figure 3.**
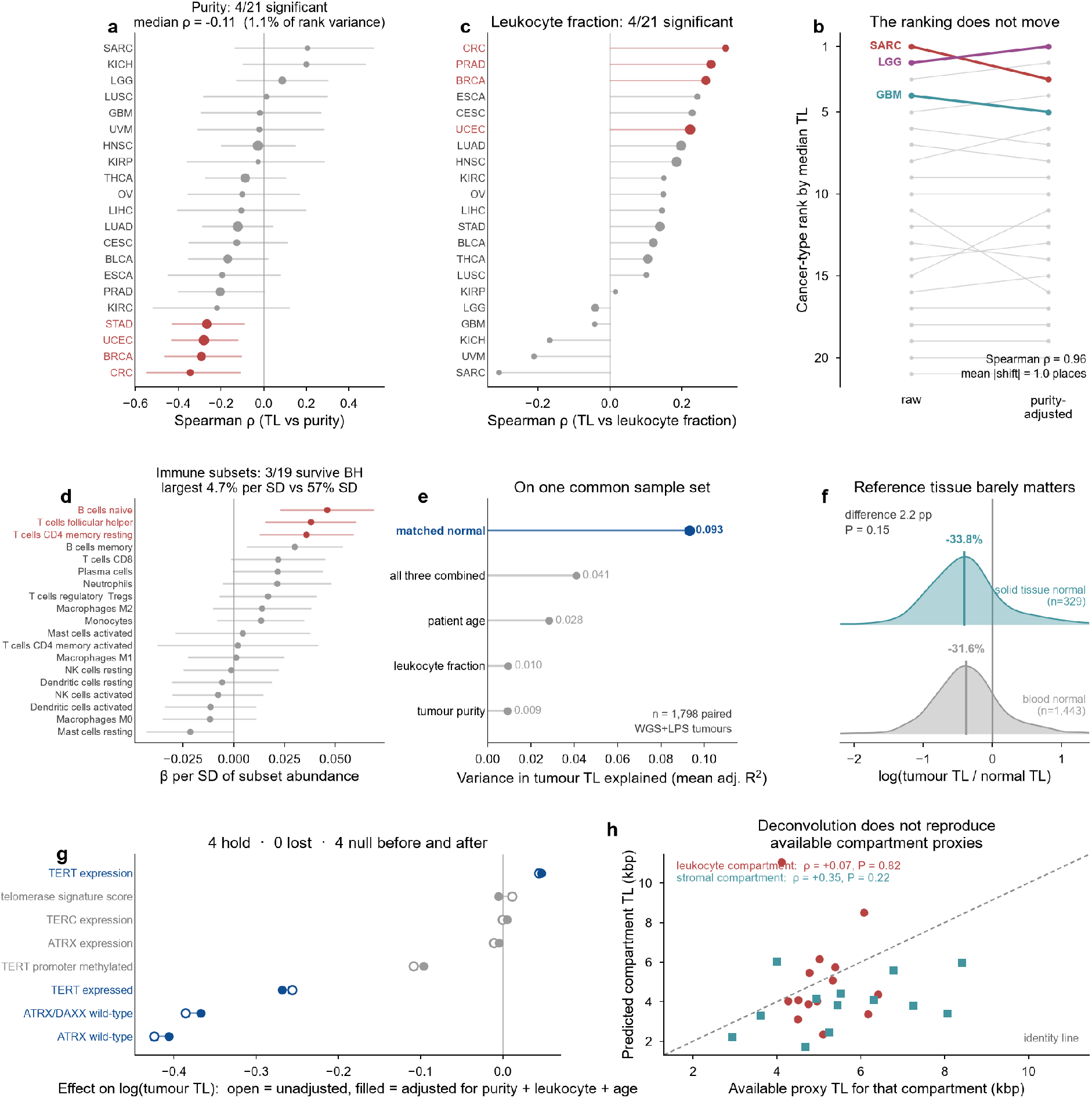
Purity, immune composition and reference tissue do not explain the pan-cancer telomere landscape. (**a**) Per-cohort correlation of telomere length with tumour purity; red marks cohorts significant after Benjamini–Hochberg correction. (**b**) Cancer-type ranking by median telomere length, raw versus purity-adjusted marginal means. (**c**) Per-cohort correlation with methylation-derived leukocyte fraction. (**d**) Standardised effect of each immune subset’s absolute abundance on telomere length, adjusted for purity and age. (**e**) Variance explained by each predictor, all computed on one common sample set. (**f**) Distribution of log(tumour/normal) telomere length by which normal tissue was used. (**g**) Published associations before (open) and after (filled) adjustment for purity, leukocyte fraction and age. (**h**) Model-predicted compartment telomere lengths against the available external proxy measurements for those compartments in the same cohorts, with the identity line. Neither proxy is ground truth.

### Immune and stromal composition

Leukocyte fraction, estimated from DNA methylation and therefore independent of both the copy-number purity call and expression-based deconvolution [34], is positively associated with telomere length in 16 of 21 cohorts (sign test *P* = 0.027) but is strongly collinear with purity (ρ = −0.73) and does not survive full adjustment (Fig. 3c). Of 19 testable immune subsets quantified by expression deconvolution [35], three survive multiple-testing correction — naive B cells (β = +0.046 per SD, *q* = 0.0014), follicular helper T cells and resting memory CD4 T cells — all positive, which is consistent with naive lymphocytes carrying the longest telomeres of circulating leukocytes. The magnitudes are nonetheless trivial: the largest is 4.6% per standard deviation against a within-cohort telomere-length standard deviation of 57% (Fig. 3d).

Computed on one common sample set (n = 1,798 paired WGS and LPS tumours), the comparison is unambiguous: matched-normal telomere length explains a mean adjusted *R*² of 0.093, against 0.041 for purity, leukocyte fraction and age combined, 0.028 for age alone, 0.010 for leukocyte fraction and 0.009 for purity (Fig. 3e).

### Reference tissue

Barthel preferred blood over solid tissue as the matched normal wherever both were available, so 80.7% of the 8,953 pairs are tumour-versus-blood (Supplementary Fig. S1c). Since blood and solid tissue differ in telomere length, this could bias the tumour-versus-normal comparison. Solid normals are indeed longer than blood normals (+16.3%, *P* = 1.1 × 10⁻¹¹), but the apparent shortening is barely affected: −31.6% using blood versus −33.8% using solid tissue, a difference of 2.2 percentage points (*P* = 0.15; Fig. 3f). The bias largely cancels because tumours paired with solid normals also have longer telomeres. Barthel’s first headline claim is therefore also robust.

### Published associations

Re-testing the associations reported for this resource with and without adjustment for purity, leukocyte fraction and age, four hold and none is lost: ATRX/DAXX status (β −0.386 → −0.368), ATRX status (−0.424 → −0.407), TERT expression status (−0.256 → −0.269) and continuous TERT expression (+0.044 → +0.047), with attenuation of 4–6%. Four further associations were null both before and after (Fig. 3g).

### Composition-based deconvolution does not reproduce available compartment proxies

A mixing model fitted to bulk data yields compartment-specific telomere lengths by extrapolation, and these can be checked against external proxy measurements. Neither proxy is ground truth — blood-normal telomere length is not a direct measurement of tumour-infiltrating leukocytes, and adjacent normal tissue is not a direct measurement of tumour stroma — but a working model should at least track them. The model’s estimate of pure-leukocyte telomere length should track the measured blood-normal telomere length of the same cohort; it does not (ρ = +0.07, *P* = 0.82), and the stromal estimate predicts measured normal-tissue telomere length worse (ρ = +0.35, *P* = 0.22) than the raw tumour median does (ρ = +0.65, *P* = 0.011) (Fig. 3h). Compartment-specific telomere length is not recoverable from bulk data by composition covariates, at least by this route. This is not a deficiency of the particular deconvolution used to supply the composition covariates: expression-based methods that estimate absolute rather than relative abundance give the same compositional picture [40], and the failure lies in the extrapolation to compartment telomere length, not in the mixture weights.

### The patient-specific component, and why the between-cohort coefficient is not interpretable

The host term can be split. Following the within/between formulation of Mundlak [41], matched-normal telomere length was separated into its cohort mean and the within-cohort deviation, and both entered a single model (Fig. 1g, Fig. 5). The within-cohort coefficient — the patient’s own deviation from their cohort — is β_within = 0.417 (95% CI 0.348–0.487) in TCGA and 0.671 (0.612–0.729) in PCAWG. It is positive in every one of the 23 TCGA and 24 PCAWG cohorts, with medians of +0.46 and +0.51 (Fig. 5b, 5c), and it moves only between 0.397 and 0.438 under adjustment for purity, leukocyte fraction, age, ploidy and coverage (Fig. 5f).

The between-cohort coefficient appears at first to carry a much stronger claim: β_between = 0.942 (0.810–1.074) in TCGA and 0.933 (0.836–1.029) in PCAWG, neither interval excluding a coefficient of one. It is tempting to read this as the tissue of origin passing its telomere setpoint into the tumour intact. That reading does not survive testing, and the test is worth setting out because the failure is instructive.

### The between-cohort coefficient is an artefact of the reference tissue

In 80.7% of pairs the matched normal is blood, so the cohort mean being used as a “tissue of origin” term is a cohort mean of *leukocyte* telomere length, and the tissue interpretation is an inference. In the tumour-versus-solid-normal pairs the reference is an organ-matched non-neoplastic tissue. That is much closer to the tissue of origin than blood is, though it is still not a direct measurement of the progenitor cell’s telomere setpoint. Refitting the decomposition separately by matched-normal type splits the estimate apart (Supplementary Fig. S5, Supplementary Table S16). With an **organ-matched solid-tissue normal** as the reference, β_between = 0.442 (0.242–0.641), which excludes a coefficient of one decisively (*P* = 7.9 × 10⁻⁸). With a **blood-normal reference**, β_between = 1.219 (1.035–1.404). The two differ at z = −4.90, *P* = 9.7 × 10⁻⁷.

Cohort composition does not explain the gap. The solid-normal arm lacks lower-grade glioma and sarcoma, the two long-telomere outliers that dominate the cross-cancer ranking; restricting the blood arm to the solid arm’s own ten cohorts leaves β_between at 1.272 (1.005–1.539), essentially unchanged.

The explanation is arithmetic. An ordinary-least-squares slope scales inversely with the spread of its predictor. Cohort-mean blood telomere length varies little between cancer cohorts, because blood is blood (between-cohort SD 0.133 on the log scale); cohort-mean tumour telomere length varies a great deal (SD 0.194). Regressing the second on the first therefore returns a slope near one for reasons of scale. Cohort-mean *tissue* telomere length varies far more (SD 0.291), and the slope falls accordingly. The ratio of the size-weighted between-cohort coefficients across the arms is 2.88 against a predictor-spread ratio of 2.19. The agreement is closer still for the unweighted cohort-level slope, for which the identity slope = *r* × SD(tumour) / SD(normal) holds exactly: there the ratio is 2.28 against the same spread ratio of 2.19, and all three arms lie within the band that identity allows (Supplementary Fig. S5d). Meanwhile the between-cohort *correlation*, which is scale-free, is stable throughout (*r* = +0.66 solid, +0.58 blood, +0.75 blood restricted). The association between a cohort’s normal and tumour telomere length is real and consistent. The coefficient of one is not a finding.

Two consequences follow, and both narrow the claim. First, no statement about one-to-one tissue-of-origin correspondence is made in this work. Second, in the solid-normal arm the within and between coefficients are indistinguishable (0.420 versus 0.442, *P* = 0.88), so the two-level separation seen in the pooled analysis is itself a property of using blood as the reference and is not reproduced when an organ-matched solid-tissue reference is used. PCAWG cannot arbitrate: its source table does not record control tissue type, and its controls are predominantly blood, so β_between = 0.933 carries the same ambiguity untestably.

The individual-level coefficient is also not demographic structure in disguise. Sex is a classical determinant of leukocyte telomere length and is one here: men have shorter matched-normal telomeres than women after adjustment for age and cancer type (β = −0.047, *P* = 0.013). Adding it changes nothing. Across a nested ladder running from cancer type alone through age, sex, purity, leukocyte fraction, ploidy, coverage and finally continental admixture proportions, the coefficient on the patient’s own normal telomere length moves only between 0.406 and 0.429 — a total spread of 0.023, every rung fitted on one common complete-case set of 1,710 pairs (Fig. 5e, Supplementary Table S19). The patient-specific term is therefore not a re-expression of age, sex or continental ancestry.

What survives this is the part that matters. β_within is 0.420 with solid normals, 0.454 with blood normals and 0.499 with blood normals restricted to the solid arm’s cohorts — stable whichever tissue supplies the reference, because it is a within-cohort quantity and does not depend on between-cohort spread. The patient-specific component is the robust finding; the between-cohort term is real as an association but its magnitude is not identified by these data.

The magnitude should not be overstated either. Partitioning the variance of tumour telomere length gives 6.1% individual, 8.6% cohort-level and 85.3% residual in TCGA, and 15.7%, 11.1% and 73.2% in PCAWG (Fig. 5g, 5h); the cohort-level shares inherit the interpretive problem above and are reported for completeness rather than as a decomposition of mechanism. The claim is that a patient’s own telomere length is a measurable, replicated component of their tumour’s, large enough to matter and so far unmodelled — not that it dominates.

### A germline variable predicts the host term, but the tumour-side test is underpowered

If the matched-patient association reflects constitutional telomere biology, a variable fixed at conception should predict it. Genetic ancestry is such a variable: it cannot be altered by the tumour, and African ancestry is robustly associated with longer leukocyte telomeres. Continental admixture proportions were obtained for 1,314 of the blood-normal pairs [42] and used as a germline proxy — not as a polygenic score for telomere length, which would require controlled-access genotypes.

The germline proxy recovers the expected normal-telomere association. African ancestry proportion predicts the patient’s own normal telomere length (β = +0.140, *P* = 6.4 × 10⁻⁴, adjusted for cancer type; +15.1% across the full ancestry range), recovering the established association within this resource and confirming that the normal telomere measurement used throughout carries genuine constitutional signal.

The tumour-side test is inconclusive, and structurally so. Ancestry does not significantly predict tumour telomere length (β = +0.038, 95% CI −0.082 to +0.159, *P* = 0.53). But the effect the constitutional model predicts is the within-cancer-type normal→tumour coefficient estimated on this same subset (0.468) multiplied by the ancestry effect on normal telomere length (0.140), which is +0.066 — and the available standard error is 0.061, larger than the effect itself. Power to detect it is 19%, and roughly 9,000 patients would be needed for 80%; this resource provides 1,314. Treating ancestry as an instrument gives +0.274 with a 95% interval of −0.597 to +1.145, which contains the ordinary-least-squares estimate, zero and one alike. First-stage strength (*F* = 11.7) is not instrument validity, and continental ancestry cannot reasonably be assumed to satisfy the exclusion restriction — it is associated with a great many genetic, environmental and social variables that could act on tumour telomere length independently of the patient’s own telomere length. That estimate is therefore reported as an exploratory sensitivity analysis and is not interpreted causally. This is not evidence against a germline contribution reaching the tumour. It is a demonstration that the question cannot be settled at this scale, together with an estimate of the scale required.

### Sensitivity to the purity estimator, and a demonstration of circularity

The choice of purity estimator is not neutral, and the data show why (Supplementary Fig. S2). Consensus Purity Estimate (CPE) combines four measures [32], one of which — LUMP — is itself derived from leukocyte-specific unmethylation. Correlating each estimator against the methylation-derived leukocyte fraction used as a predictor elsewhere in this work gives LUMP ρ = −0.87, CPE ρ = −0.83, ESTIMATE ρ = −0.75 [33], ABSOLUTE ρ = −0.73 [38] and immunohistochemistry ρ = −0.28. Copy-number-derived ABSOLUTE is the least entangled of the genomic estimators, which is why it is used as primary throughout.

This has a measurable consequence: the apparent purity effect on telomere length roughly doubles with the more entangled estimators (pooled β = −0.55 for ESTIMATE, −0.50 for CPE, −0.39 for LUMP, −0.25 for ABSOLUTE, −0.20 and non-significant for immunohistochemistry). The ordering is exactly what shared measurement input predicts. No conclusion changes, however: ABSOLUTE and CPE agree at ρ = 0.66 overall and ρ = 0.85 across per-cohort effects, and the cross-cancer ranking is robust under either (ρ = 0.97 and 0.91 against the raw ranking). The host effect survives conditioning on either purity estimator and on stage, ploidy, genome doublings and subclonal fraction, individually and all together (β = 0.434 unadjusted, 0.464 with all five entered simultaneously). This conditioning set differs from the one in Fig. 4f, which substitutes leukocyte fraction, age and coverage for stage and subclonal fraction and ends at β = 0.390; the host term is stable in the 0.39–0.48 range under every combination tried.

**Figure 4.**
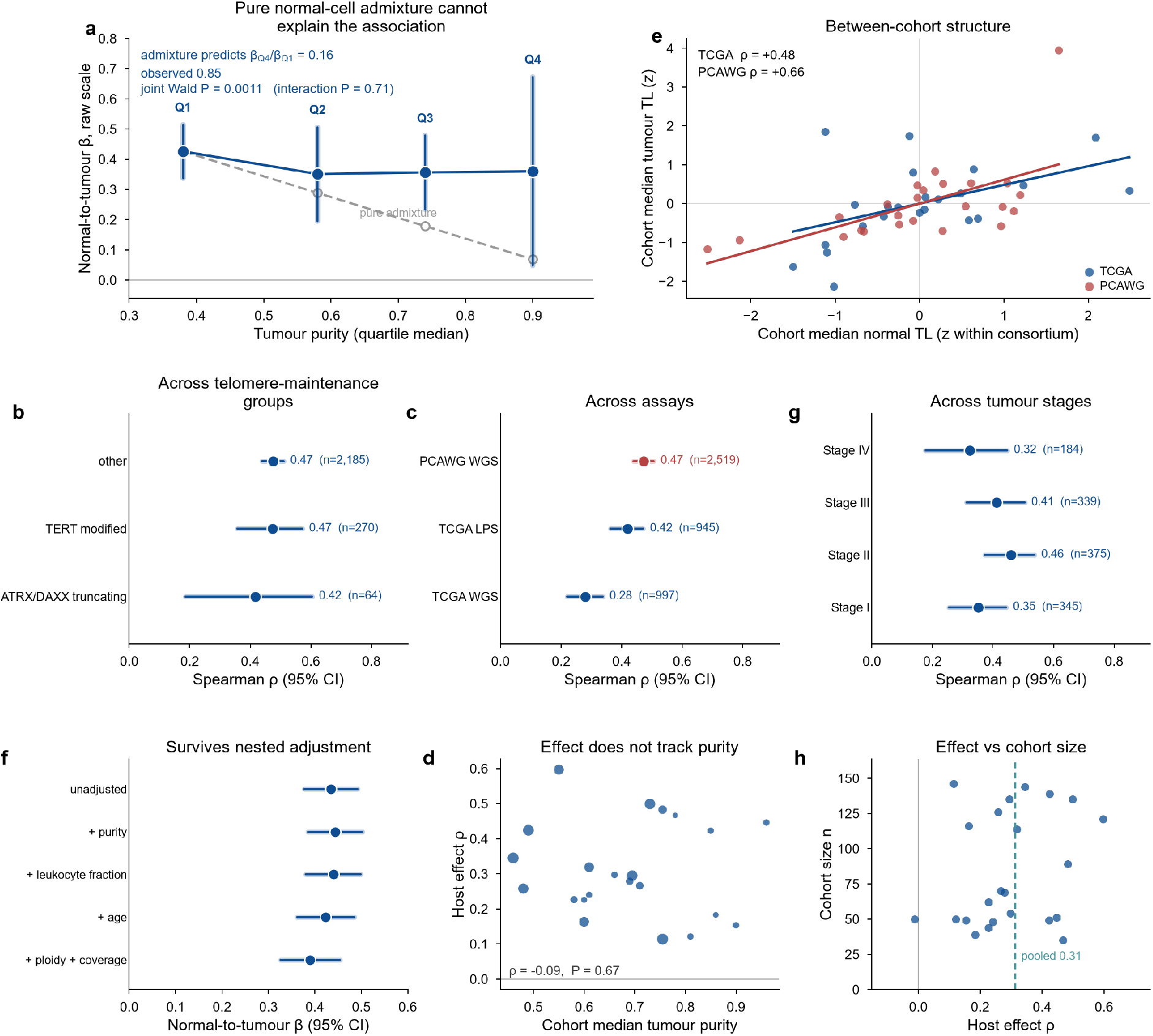
Pure normal-cell admixture cannot explain the association, which is observed across mechanism, assay and stage. (**a**) Normal→tumour coefficient by tumour purity quartile, fitted on the raw telomere-length scale because the admixture identity is a raw-scale statement, with the trajectory predicted under pure admixture (grey dashed). (**b**) By telomere maintenance mechanism (PCAWG). (**c**) By assay and cohort. (**d**) Per-cohort effect size against cohort median purity. (**e**) Cohort mean normal and tumour telomere length, z-scored separately within each consortium. (**f**) Nested adjustment of the normal→tumour coefficient. (**g**) By tumour stage. (**h**) Effect size against cohort size.

**Figure 5.**
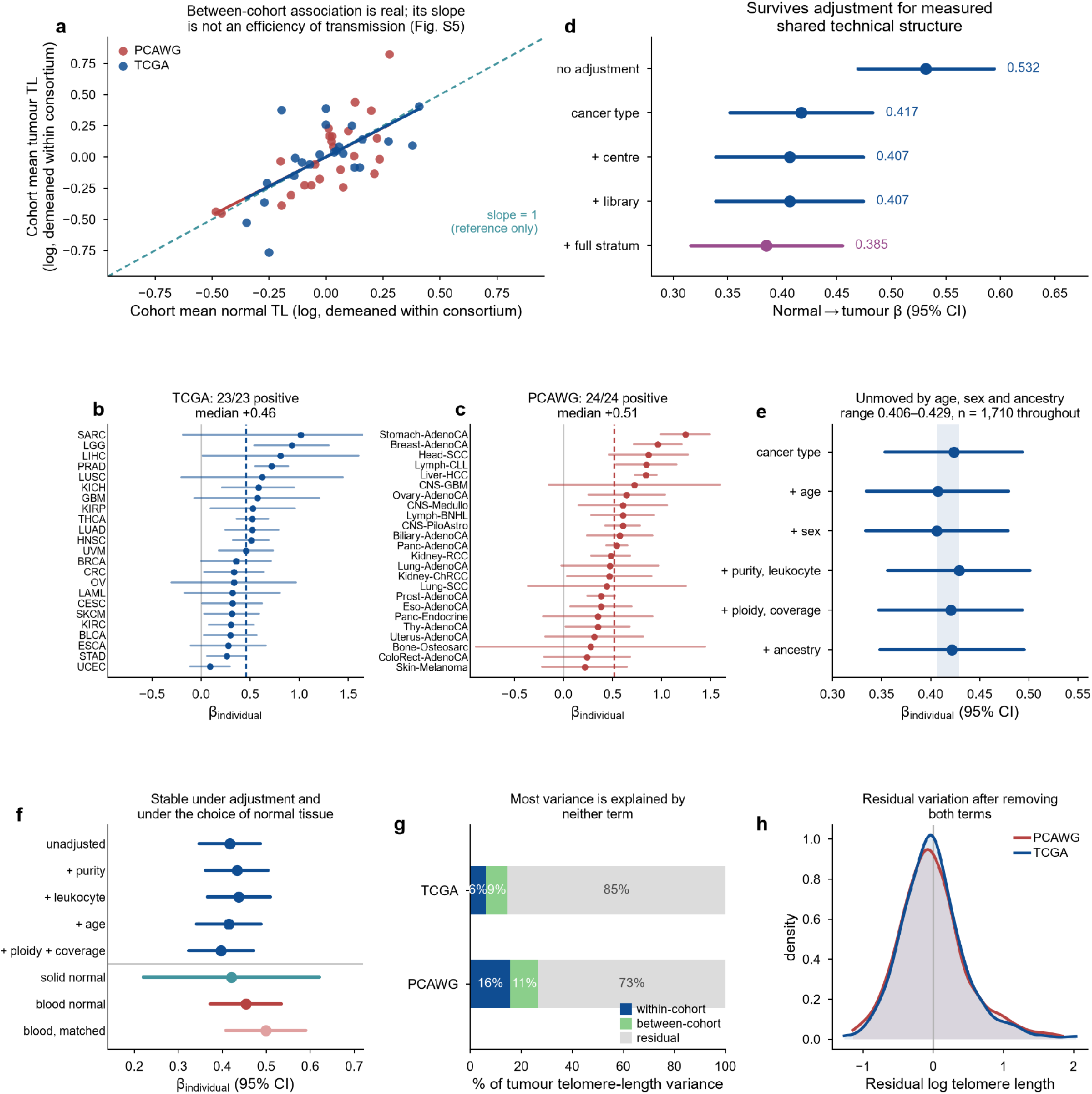
The patient-specific coefficient, and what it survives. (**a**) Cohort mean normal against cohort mean tumour telomere length, demeaned separately within each consortium, with the fitted between-cohort coefficients. The dashed line marks a slope of one and is a reference only: as shown in Supplementary Fig. S5, this slope is not an efficiency of transmission. (**b**, **c**) Individual-level coefficient in each TCGA and PCAWG cohort, with 95% confidence intervals; the dashed line is the median. (**d**) The normal→tumour coefficient with technical structure entered as fixed effects, ending with one fixed effect for the complete cancer-type × sequencing-centre × library-type stratum — the same unit the permutation test shuffles within. (**e**) The individual-level coefficient under nested demographic and compositional adjustment, every rung fitted on one common complete-case set so that the sample cannot change under the coefficient; the shaded band spans the full range across the ladder. (**f**) The individual-level coefficient under every adjustment tested and under each choice of matched normal tissue; the rule separates adjustment models above from reference-tissue arms below. (**g**) Variance of tumour telomere length partitioned into within-cohort, between-cohort and residual components. The between-cohort share is not a tissue-of-origin contribution (Supplementary Fig. S5). (**h**) Residual variation after removing both terms.

### Analyses that did not support a claim

Three lines of investigation are reported because they were run, not because they worked.

#### An apparent age artefact in glioma, which did not survive its control (Supplementary Fig. S3)

Since telomere length falls with age and cohorts differ in median age by up to 26 years, cohort age is a candidate explanation for the cross-cancer ranking. It is not one in general (cohort median age versus cohort median telomere length ρ = −0.16, *P* = 0.50). Lower-grade glioma, however, is a marked age outlier (median 42 years against 61 elsewhere) with the steepest within-cohort age slope, and restricting all cohorts to ages 50–70 moved it from rank 2 to rank 14 while leaving sarcoma at rank 1 and glioblastoma at rank 3. The interpretation did not survive conditioning on *IDH* status. IDH-mutant and IDH-wild-type lower-grade gliomas differ in both age (median 39 versus 52, *P* = 0.008) and telomere length (5.95 versus 3.09 kbp, *P* = 0.003) [43,44,45]; within IDH-mutant gliomas alone the age slope is not significant (β = −0.093 per decade, *P* = 0.10); age-matching silently shifts the IDH-mutant share from 76% to 58% (Fisher *P* = 0.004); and IDH-mutant gliomas restricted to ages 50–70 return to rank 1 (median 5.25 kbp, *P* = 0.007 against all other cohorts). Holding molecular subtype constant restores the original finding. The apparent glioma age effect therefore reflects *IDH*-defined molecular composition rather than age itself, consistent with differences in telomere-maintenance biology among glioma subtypes [46]; the analysis does not establish that alternative lengthening of telomeres specifically drives the ranking, since *IDH*-mutant glioma is not synonymous with ALT. Barthel’s claim stands.

#### Cell lines cannot cross-validate the ranking (Supplementary Fig. S4a–e)

Cancer cell lines contain no host, and so appeared to offer a decisive test. They do not, for structural reasons. The correlation between CCLE lineage medians and TCGA cancer-type medians is ρ = −0.315 with a bootstrap 95% confidence interval of [−0.62, +0.27] and 20% of resamples positive — the test cannot distinguish the hypotheses. Two reasons: lineage medians are unstable, with a median bootstrap interval width of 126% of the entire between-lineage spread; and culture selection is total, with 0 of 19 CCLE lower-grade glioma lines and 0 of 45 glioblastoma lines carrying an *IDH1* mutation against 76% of TCGA lower-grade gliomas. Consistent with immortalisation erasing the host term, donor age does not predict cell-line telomere content (ρ = −0.06, *P* = 0.30).

#### Telomere length has no prognostic value here (Supplementary Fig. S4f–h)

Using the curated TCGA clinical endpoints [47] with progression-free interval as the primary outcome, neither tumour telomere length (HR 0.97, *P* = 0.47), host normal telomere length (HR 0.95, *P* = 0.16) nor the tumour/normal ratio (HR 1.01, *P* = 0.85) was associated with outcome; overall survival gave the same picture, and adjustment for age and purity changed nothing. Discrimination was at chance in every form: mean concordance 0.498 for tumour telomere length, 0.500 for the ratio and 0.482 for host telomere length across 13 cancer types. With ∼1,900 patients and ∼650–690 events this is a well-powered null for the pooled question, not an absence of power. It is not, however, a test of every disease: prostate cancer — the one setting in which host telomere length has been reported to be prognostic [36] — contributes too few progression events to enter the per-cohort discrimination analysis at all.

## Discussion

The pan-cancer telomere landscape has been read as a statement about cancers. The results here suggest it is partly a statement about patients.

Three findings support that reading. Tumour telomere length correlates with the patient’s own normal telomere length in essentially every cohort of two consortia measured with two different estimators, at a magnitude that survives a stringent within-stratum permutation test. The individual-level coefficient is stable at 0.42–0.50 across cohorts, across adjustment, and across the choice of reference tissue. And matched-normal telomere length alone explains variance comparable to cancer type, although its unique individual-level component is smaller.

The individual-level coefficient is the interpretable result, and it is interpretable precisely because it does not depend on between-cohort spread. It says that among patients with the same cancer, one whose constitutional telomeres are 20% longer than their cohort’s average will have a tumour roughly 8–13% longer. Demanelis and colleagues showed that telomere length varies systematically between normal tissues while remaining correlated within individuals [29]; the present result extends the within-individual half of that observation into malignancy.

The tissue-level half is where this work initially over-read its own data, and the correction is worth stating plainly because the same trap is available to anyone using this resource. A between-cohort regression coefficient is not an efficiency of transmission. Its magnitude depends on how much the predictor varies across the units being compared, and when the predictor is leukocyte telomere length — which barely differs between cancer cohorts — a slope near one carries no biological content. The scale-free correlation is stable and the association is genuine, so cohort-level telomere length and tumour telomere length do travel together; the parsimonious reframing of Barthel’s second claim, that sarcoma and glioma may have long telomeres partly because their tissues of origin do, remains available and is consistent with cell of origin dominating molecular classification more generally [48]. But this work cannot say how much of the tissue setpoint is carried through transformation, and the solid-normal arm provides no evidence that the within- and between-cohort coefficients differ.

The individual-level term connects to a large epidemiological literature. Leukocyte telomere length is heritable [26], has a mapped common-variant architecture [27], and genetically-predicted longer telomeres are associated with the risk of several cancers [28]. Under a causal constitutional-transmission model, germline determinants of telomere length would be expected to contribute to tumour telomere length. That model is not established here, but it yields a clean testable prediction: a telomere-length polygenic score should associate with tumour telomere length, and the constitutional axis would then be a candidate mediator of some germline telomere–cancer associations. Testing that requires individual-level germline genotypes under controlled access and remains the clearest next experiment. The ancestry analysis above bounds what it would take: with an effect of the size the constitutional model predicts, roughly 9,000 tumour/normal pairs on a calibrated assay would be needed for 80% power, against the 1,314 available here.

It is worth being equally clear about what these results do not show. The individual-level term explains 6.1% of tumour telomere-length variance in TCGA and 15.7% in PCAWG, and even adding the between-cohort share — which, as shown above, is not identified as a tissue contribution — reaches only 14.7% and 26.8%. Most of the variance remains unexplained by the individual- and between-cohort terms — residual variance contains tumour biology, measurement error, unmeasured host factors and unmodelled technical variation alike — and the mechanisms established for tumour-intrinsic variation — telomerase reactivation [7], *TERT* promoter mutation [8,9,10], ALT following *ATRX*/*DAXX* loss [11,12,13] — are unaffected. Indeed all four published telomere associations testable in this resource survive adjustment essentially unchanged. Nor is causal direction established: matched normal and tumour are measured in the same person at the same time, and shared germline or environmental determinants could produce the correlation without telomere length being carried through transformation. The matched normals are also ∼80% blood, so “constitutional” is an inference from leukocyte telomere length rather than a direct measurement of the tissue of origin in each patient.

The negative results deserve emphasis rather than burial, since they are what leaves the patient-specific association robust to the technical and compositional alternatives tested here. Tumour purity has a real but small within-cohort effect on telomere length and essentially no effect on the cross-cancer ranking. Immune composition contributes at the ∼1% level despite a mechanistically plausible route — telomere length regulates *IL1R1* through TRF2 and thereby modulates immune signalling in the tumour microenvironment [15], a relationship that operates in the opposite causal direction to the one tested here and would not be visible as a compositional association. Reference-tissue choice changes the tumour-versus-normal gap by 2.2 percentage points. And the mixing-model approach to recovering compartment-specific telomere lengths, which motivated much of this work, fails against the available external proxies. That last result is worth stating plainly for anyone contemplating telomere deconvolution from bulk data: when the model’s leukocyte estimate is checked against measured blood-normal telomere length from the same cohorts, it does not correlate with it at all. The prognostic null belongs in the same list: leukocyte telomere length is associated with all-cause mortality in general populations [49], but neither tumour nor host telomere length discriminated outcome across the thirteen cancer types testable here.

That null needs reconciling with the one published report that looked. Livingstone and colleagues found no association between tumour telomere length and biochemical relapse in localised prostate cancer — which agrees with the result here — but did find one for the patient’s non-tumour telomere length (HR 0.77 per unit, *P* = 0.014 as a continuous variable; HR 2.02, *P* = 0.0016 at an outcome-optimised 3.9 kbp cut point, n = 290), and interpreted it as reflecting host factors influencing patient risk [36]. Their interpretation anticipates the argument made here. The disagreement is narrower than it appears: prostate cancer is absent from the discrimination analysis in this work because it does not reach the minimum of fifteen progression events, so the two analyses do not test the same disease, the same endpoint, or the same predictor form. The honest reading is that host telomere length may well be prognostic in specific settings and that this work is not powered to detect it disease by disease; what it does establish is that neither telomere measure discriminates across the thirteen cancer types where the test could be run.

The calibration finding stands somewhat apart and may be the most immediately actionable. That whole-exome telomere estimates fail to recover age-related attrition in blood is not a subtle bias; it is a failure of the measurement to track the single best-established quantitative fact about telomeres in human tissue. Since exome data constitute 78.6% of the most widely used pan-cancer telomere resource, and since the WXS-derived cross-cancer ranking does not agree with the WGS-derived one, any analysis of that resource should either restrict to whole-genome and low-pass data or explicitly justify why exome estimates are fit for its purpose. The failure was not reproduced in the small CCLE comparison, which suggests it is context-dependent rather than an inevitable consequence of exome capture. Whether that dependence reflects cellular admixture, specimen or DNA quality, technical batch, capture protocol or some combination cannot be resolved here, and the 36-line comparison establishes only that the discordance is not universal to exome sequencing.

A route past both the calibration and the deconvolution failure exists in principle. Telomere length would need to be measured in the compartments directly rather than inferred from a mixture, which single-cell telomere quantification and quantitative fluorescence in situ hybridisation on tissue sections make conceivable [50,51] but which no current pan-cancer resource provides at the scale required. Until such data exist, the host term is best handled as a covariate rather than removed.

Several limitations follow from the design. All analyses are re-analyses of publicly released per-sample derived measurements and therefore inherit the biases of the original measurement pipelines. The two estimators differ (TelSeq for TCGA, TelomereHunter for PCAWG), which is a strength for replication and a weakness for pooling. TCGA and PCAWG share donors, addressed here by a restricted-cohort sensitivity analysis rather than by exclusion at the donor level. Most consequentially, 80.7% of matched normals are blood rather than the tissue of origin, which is why the between-cohort coefficient is not identified here; only 300 pairs across 12 cohorts have a solid-tissue normal, and no patient in the paired set has both, so the within-patient contest that would settle it cannot be run. The two independent compartment estimates used — copy-number purity and methylation-derived leukocyte fraction — are mutually inconsistent for 7.7% of samples, in the sense that they sum to more than one. Coverage is unavailable for exome samples in the source table, so the coverage covariate exists only for the whole-genome and low-pass set. And the analysis set is modest by pan-cancer standards: 1,706 primary tumours, of which the two cohorts most central to the headline claims contain 40 and 89.

The practical recommendations are narrow and, I think, defensible. Studies that use bulk tumour telomere length as a variable should carry matched normal telomere length as a covariate where it exists, and should say so where it does not. Cross-cancer comparisons of telomere length should be read as comparisons of patient populations and tissues as much as of cancers. And telomere estimates from exome sequencing of clinical specimens should not be used as a quantitative phenotype without local calibration against a known biological gradient — age is a convenient one, and blood normals make it easy.

More broadly, the result is an instance of a general problem in cancer genomics: a measurement taken from a tumour is not automatically a measurement of the tumour [52]. Bulk telomere length has been treated as a cancer phenotype for a decade. It is better understood as a composite phenotype containing a replicated patient-specific component, measurable and so far unmodelled, alongside cohort- and tissue-associated variation whose magnitude these data cannot resolve.

## Methods

### Data sources

#### Telomere length, TCGA

Per-sample telomere length estimates for 18,430 TCGA samples were obtained from Supplementary Table 1 of Barthel et al. [17]. Telomere length was estimated in that work with TelSeq (*k* = 7) as a read-group-weighted mean and is reported in kilobase pairs [20]. The table provides an “Unpaired Set” (18,430 samples; one row per sample) and a “Paired Set” (8,953 tumour/normal pairs with normal telomere length, tumour telomere length, the log ratio, and genomic annotations including *TERT* expression status, *ATRX*/*DAXX* status, *TP53* mutation and telomerase signature score).

#### Telomere content, PCAWG

Matched tumour and control telomere content for 2,519 PCAWG specimens across 35 histologies was obtained from Supplementary Data 1 of Sieverling et al. [19], where it was estimated with TelomereHunter [21]. The 2022 Author Correction to that paper was checked and concerns only the placement of the consortium author list; no data, figures or supplementary files were altered.

#### Tumour purity and ploidy

ABSOLUTE purity, ploidy, genome doublings and subclonal genome fraction were taken from the PanCanAtlas TCGA_mastercalls.abs_tables_JSedit.fixed.txt release (10,786 rows), derived by the ABSOLUTE method [38]. Consensus Purity Estimate and its four components (ESTIMATE, ABSOLUTE, LUMP, immunohistochemistry) were taken from Supplementary Data 1 of Aran et al. [32]; the ESTIMATE component derives from the method of Yoshihara et al. [33].

#### Immune composition

Leukocyte fraction (DNA-methylation-derived) and CIBERSORT relative immune-cell fractions [35] for 22 cell types were taken from the PanCanAtlas immune-landscape release accompanying Thorsson et al. [34]. Absolute subset abundance was derived as leukocyte fraction × relative fraction, since CIBERSORT relative fractions sum to one within the leukocyte compartment.

#### Clinical outcomes

Overall survival, disease-specific survival, disease-free interval and progression-free interval were taken from the TCGA Clinical Data Resource [47]. Progression-free interval was used as the primary endpoint and overall survival as secondary, following that resource’s own usage guidance. Redacted cases were excluded.

#### Cell lines

Telomere content for CCLE cell lines (TelSeq, WGS and WES) with lineage, donor age and *IDH1* mutation annotation was taken from Supplementary File 1 of Hu et al. [37].

#### IDH status

*IDH1* and *IDH2* mutation calls for TCGA lower-grade glioma and glioblastoma were retrieved from cBioPortal PanCanAtlas studies [53,54]. The _sequenced sample list was used as the denominator, so that absence of a mutation call is interpreted as wild-type only for samples actually sequenced. Silent and synonymous variants were excluded.

### Harmonisation

Barthel sample identifiers are 15-character TCGA sample barcodes (verified for 18,430 of 18,430 rows) and were used as the merge key throughout. ABSOLUTE calls are reported at the same level. Leukocyte fraction and CIBERSORT are reported per aliquot (28-character barcodes); these were truncated to 15 characters and collapsed to one value per sample by median. CPE identifiers are 16-character barcodes including the vial letter and were truncated to 15 characters, again collapsing duplicates by median. TCGA sample-type codes were retained as supplied: TP (primary tumour), NB (blood-derived normal), NT (solid tissue normal), TM (metastatic tumour), TB (blood-derived tumour). Merge yields are reported in full in Supplementary Fig. S1 and were 96.2% (ABSOLUTE), 96.7% (leukocyte fraction) and 93.5% (CIBERSORT) for primary tumours; all three resources are tumour-only by construction, so normals match at 0%.

### Analysis set

Analyses were restricted to WGS and LPS libraries on the basis of the calibration result (Fig. 2), to primary tumours (sample type TP) for cross-cancer analyses, and to cancer types with at least 25 qualifying samples. The threshold was varied from 10 to 80 as a sensitivity analysis (Supplementary Fig. S5h). Telomere length was log-transformed for linear modelling because of right skew, except for the admixture falsification, which was fitted on the raw scale because the two-compartment mixture identity is defined on that scale.

### Statistical analysis

Within-cohort associations were assessed by Spearman rank correlation with percentile bootstrap confidence intervals (2,000–5,000 resamples of observed pairs). Linear models were fitted per cancer type and pooled by inverse-variance meta-analysis; this was preferred to a mixed model because maximum-likelihood mixed-model fits failed to converge on several specifications. Heterogeneity is reported as *I*². Multiple testing across cancer types or immune subsets was controlled by the Benjamini– Hochberg procedure [39] at *q* < 0.05.

### Permutation test

To separate the matched-patient association from batch structure, matched normals were permuted among patients within strata defined by cancer type × sequencing centre × library type (TCGA; 40 strata with ≥ 8 pairs) or by histology (PCAWG; 25 strata), preserving cancer type, sequencing centre and library type while destroying the pairing. 10,000 permutations were used; the observed statistic is reported as a *z* score against the permutation null.

### Within/between decomposition

Matched-normal log telomere length was split into its cohort mean (between-cohort) and the deviation from that mean (within-cohort, interpreted as individual constitution), and both terms entered a single model, following Mundlak [41]. The between-cohort term is *not* interpreted as an efficiency of tissue-of-origin transmission; the analysis by matched-normal type showing why is described below. Equality of the two coefficients was tested by a linear contrast, and the between-cohort coefficient was additionally tested against unity. Variance shares were computed as the variance of each fitted component divided by the variance of the outcome.

### Fixed-effects adjustment

As a direct complement to the permutation test, tumour log telomere length was regressed on matched-normal log telomere length with technical structure entered as fixed effects, in a nested sequence ending with a single factor for the complete cancer-type × sequencing-centre × library-type stratum. Partial *R*² is reported against the model without the normal-telomere term, and a residual-residual Spearman correlation was computed after residualising both variables on that complete stratum.

### Sex and demographic sensitivity

Sex was taken from the source table and entered alongside age, purity, leukocyte fraction, ploidy, coverage and continental admixture proportions in a nested ladder, to test whether the individual-level coefficient is stable once demographic and ancestry structure is removed rather than whether any individual covariate is significant. Because the paired set contains metastatic and blood-derived tumours as well as primary ones, while purity, ploidy, leukocyte fraction and coverage are annotations of a specific tumour sample, these covariates were joined on the exact 15-character tumour sample barcode rather than at patient level. The nested analysis was then restricted to the 1,710 pairs with complete data on every covariate, and that identical set was used at every rung, so the coefficient cannot appear stable merely because the sample changes between models.

### Matched-normal type

The within/between decomposition was refitted separately on tumour/blood-normal and tumour/solid-normal pairs, at a common minimum cohort size of ten, and the blood arm was additionally restricted to the cohorts present in the solid arm so that the two are compared on the same cancers. Coefficients were contrasted by a two-sample z test on the difference. The cohort-level (unweighted) regression slope is reported alongside the size-weighted between-cohort coefficient, because the identity slope = *r* × SD(tumour) / SD(normal) holds exactly only for the former.

### Genetic ancestry

Continental admixture proportions were taken from the PanCanAtlas ancestry release [42], averaged to one value per patient, and merged to the blood-normal pairs. African ancestry proportion was regressed on log telomere length with a cancer-type term, first in the matched normal as a validity check against the established association and then in the tumour. An instrumental-variable estimate used ancestry as the instrument for constitutional telomere length, with the first-stage *F* statistic reported. Power was computed against the effect implied by the observed individual-level coefficient.

### Admixture test

Because the mixture identity Y = *p*T + (1 − *p*)H is defined on raw telomere length, this analysis alone was fitted on the raw scale. Samples were divided into purity quartiles and the coefficient estimated within each; the ratio predicted between the highest and lowest quartile was computed from their median purities. That quartile comparison is a visualisation, since purity varies within a quartile and the median-based ratio is therefore approximate.

The formal test is the continuous model Y = α + β_H·H + β_p·*p* + β_H×p·H·*p* + ε, fitted on all samples with a purity call. Under pure admixture with the tumour component conditionally independent of host telomere length given purity, E[Y | H, *p*] = *p*·E[T | *p*] + (1 − *p*)·H, so β_H = 1 and β_H×p = −1. Both restrictions were tested jointly by a Wald *F* test and individually by *t* contrasts. This assumes the main effect of purity adequately represents *p*·E[T | *p*]; the joint test was therefore repeated with a natural cubic spline (4 degrees of freedom) and with a quadratic in purity, keeping β_H and β_H×p linear, as a sensitivity analysis.

### Survival analysis

Cox proportional-hazards models were stratified by cancer type, with predictors standardised within cancer type. Discrimination was assessed by Harrell’s concordance index computed per cancer type in cohorts with at least 50 patients and 15 events, and compared between predictors by Wilcoxon signed-rank test.

### Software

Analyses used Python 3.13 with NumPy [55], SciPy [56], pandas, statsmodels and lifelines. Figures were produced with Matplotlib [57].

### Reproducibility

Every value reported in the text and every value plotted in a figure is written to a tab-separated file under results/ by a numbered analysis script, and figure scripts read only from those files. A provenance log is maintained alongside the analysis recording, for every downloaded input, the source URL, the publication it accompanies, the download date, the SHA-256 hash and the row count; the source URLs and accompanying publications are additionally listed under Data availability.

## Data availability

All data analysed in this study are previously published and publicly available. Telomere length estimates: Supplementary Table 1 of Barthel et al. (doi:10.1038/ng.3781). PCAWG telomere content: Supplementary Data 1 of Sieverling et al. (doi:10.1038/s41467-019-13824-9). ABSOLUTE purity and ploidy: NCI Genomic Data Commons PanCanAtlas publication page. Consensus Purity Estimate: Supplementary Data 1 of Aran et al. (doi:10.1038/ncomms9971). Leukocyte fraction and CIBERSORT estimates: PanCanAtlas immune-landscape release accompanying Thorsson et al. (doi:10.1016/j.immuni.2018.03.023). Clinical outcomes: TCGA Clinical Data Resource (doi:10.1016/j.cell.2018.02.052). CCLE telomere content: Supplementary File 1 of Hu et al. (doi:10.7554/eLife.66198). *IDH* mutation calls: cBioPortal PanCanAtlas studies. Genetic ancestry and continental admixture proportions: PanCanAtlas ancestry release accompanying Carrot-Zhang et al. (doi:10.1016/j.ccell.2020.04.012). Individual-level GTEx telomere measurements (doi:10.1126/science.aaz6876) are under controlled access via dbGaP (phs000424) and were not used.

## Code availability

Analysis code and frozen outputs supporting all results reported here are archived on Zenodo at doi:10.5281/zenodo.22215444. All source datasets are publicly available from the original repositories listed under Data availability. A fully packaged source-data reproduction release will accompany the final published version.

## Author contributions

S.D. designed the study, performed all analyses and wrote the manuscript.

## Supplementary figure legends

Full panel-by-panel legends, nineteen supplementary tables and four supplementary notes are provided in the Supplementary Information.

**Supplementary Figure S1.** Sample selection, barcode harmonisation and merge quality control.

**Supplementary Figure S2.** Purity-estimator sensitivity, conditioning, and the circularity between consensus purity estimates and methylation-derived leukocyte fraction.

**Supplementary Figure S3.** The age/*IDH* glioma analysis, shown in full as a worked negative.

**Supplementary Figure S4.** Two arms that did not support a claim: cell-line cross-validation and clinical outcome.

**Supplementary Figure S5.** Why the between-cohort coefficient is not an efficiency of transmission, and the limits of a germline test at this sample size.

## References

1. Bonnell E, Pasquier E, Wellinger RJ Telomere Replication: Solving Multiple End Replication Problems. Frontiers in cell and developmental biology 9:668171 (2021). doi:10.3389/fcell.2021.668171 PMID 33869233.

2. de Lange T Shelterin-Mediated Telomere Protection. Annual review of genetics 52:223-247 (2018). doi:10.1146/annurev-genet-032918-021921 PMID 30208292.

3. Shay JW Telomeres and aging. Current opinion in cell biology 52:1-7 (2018). doi:10.1016/j.ceb.2017.12.001 PMID 29253739.

4. Stewart SA, Weinberg RA Telomeres: cancer to human aging. Annual review of cell and developmental biology 22:531–57 (2006). doi:10.1146/annurev.cellbio.22.010305.104518 PMID 16824017.

5. Maciejowski J, de Lange T Telomeres in cancer: tumour suppression and genome instability. Nature reviews. Molecular cell biology 18:175–186 (2017). doi:10.1038/nrm.2016.171 PMID 28096526.

6. Hanahan D Hallmarks of Cancer: New Dimensions. Cancer discovery 12:31-46 (2022). doi:10.1158/2159-8290.CD-21-1059 PMID 35022204.

7. Kim NW, Piatyszek MA, Prowse KR, Harley CB, West MD, Ho PL, et al. Specific association of human telomerase activity with immortal cells and cancer. Science (New York, N.Y.) 266:2011–5 (1994). doi:10.1126/science.7605428 PMID 7605428.

8. Horn S, Figl A, Rachakonda PS, Fischer C, Sucker A, Gast A, et al. TERT promoter mutations in familial and sporadic melanoma. Science (New York, N.Y.) 339:959–61 (2013). doi:10.1126/science.1230062 PMID 23348503.

9. Huang FW, Hodis E, Xu MJ, Kryukov GV, Chin L, Garraway LA Highly recurrent TERT promoter mutations in human melanoma. Science (New York, N.Y.) 339:957–9 (2013). doi:10.1126/science.1229259 PMID 23348506.

10. Killela PJ, Reitman ZJ, Jiao Y, Bettegowda C, Agrawal N, Diaz LA Jr, et al. TERT promoter mutations occur frequently in gliomas and a subset of tumors derived from cells with low rates of self-renewal. Proceedings of the National Academy of Sciences of the United States of America 110:6021–6 (2013). doi:10.1073/pnas.1303607110 PMID 23530248.

11. Bryan TM, Englezou A, Dalla-Pozza L, Dunham MA, Reddel RR Evidence for an alternative mechanism for maintaining telomere length in human tumors and tumor-derived cell lines. Nature medicine 3:1271–4 (1997). doi:10.1038/nm1197-1271 PMID 9359704.

12. Heaphy CM, de Wilde RF, Jiao Y, Klein AP, Edil BH, Shi C, et al. Altered telomeres in tumors with ATRX and DAXX mutations. Science (New York, N.Y.) 333:425 (2011). doi:10.1126/science.1207313 PMID 21719641.

13. Lovejoy CA, Li W, Reisenweber S, Thongthip S, Bruno J, de Lange T, et al. Loss of ATRX, genome instability, and an altered DNA damage response are hallmarks of the alternative lengthening of telomeres pathway. PLoS genetics 8:e1002772 (2012). doi:10.1371/journal.pgen.1002772 PMID 22829774.

14. Sengupta A, Vinayagamurthy S, Soni D, Deb R, Mukherjee AK, Dutta S, et al. Telomeres control human telomerase (TERT) expression through non-telomeric TRF2. eLife 14 (2025). doi:10.7554/eLife.104045 PMID 41026782.

15. Mukherjee AK, Dutta S, Singh A, Sharma S, Roy SS, Sengupta A, et al. Telomere length sensitive regulation of interleukin receptor 1 type 1 (IL1R1) by the shelterin protein TRF2 modulates immune signalling in the tumour microenvironment. eLife 13 (2024). doi:10.7554/eLife.95106 PMID 39728924.

16. Okamoto K, Seimiya H Revisiting Telomere Shortening in Cancer. Cells 8 (2019). doi:10.3390/cells8020107 PMID 30709063.

17. Barthel FP, Wei W, Tang M, Martinez-Ledesma E, Hu X, Amin SB, et al. Systematic analysis of telomere length and somatic alterations in 31 cancer types. Nature genetics 49:349–357 (2017). doi:10.1038/ng.3781 PMID 28135248.

18. ICGC/TCGA Pan-Cancer Analysis of Whole Genomes Consortium Pan-cancer analysis of whole genomes. Nature 578:82-93 (2020). doi:10.1038/s41586-020-1969-6 PMID 32025007.

19. Sieverling L, Hong C, Koser SD, Ginsbach P, Kleinheinz K, Hutter B, et al. Genomic footprints of activated telomere maintenance mechanisms in cancer. Nature communications 11:733 (2020). doi:10.1038/s41467-019-13824-9 PMID 32024817.

20. Ding Z, Mangino M, Aviv A, Spector T, Durbin R, UK10K Consortium Estimating telomere length from whole genome sequence data. Nucleic acids research 42:e75 (2014). doi:10.1093/nar/gku181 PMID 24609383.

21. Feuerbach L, Sieverling L, Deeg KI, Ginsbach P, Hutter B, Buchhalter I, et al. TelomereHunter - in silico estimation of telomere content and composition from cancer genomes. BMC bioinformatics 20:272 (2019). doi:10.1186/s12859-019-2851-0 PMID 31138115.

22. Nersisyan L, Arakelyan A Computel: computation of mean telomere length from whole-genome next-generation sequencing data. PloS one 10:e0125201 (2015). doi:10.1371/journal.pone.0125201 PMID 25923330.

23. Farmery JHR, Smith ML, NIHR BioResource - Rare Diseases, Lynch AG Telomerecat: A ploidy-agnostic method for estimating telomere length from whole genome sequencing data. Scientific reports 8:1300 (2018). doi:10.1038/s41598-017-14403-y PMID 29358629.

24. Müezzinler A, Zaineddin AK, Brenner H A systematic review of leukocyte telomere length and age in adults. Ageing research reviews 12:509–19 (2013). doi:10.1016/j.arr.2013.01.003 PMID 23333817.

25. Ye Q, Apsley AT, Etzel L, Hastings WJ, Kozlosky JT, Walker C, et al. Telomere length and chronological age across the human lifespan: A systematic review and meta-analysis of 414 study samples including 743,019 individuals. Ageing research reviews 90:102031 (2023). doi:10.1016/j.arr.2023.102031 PMID 37567392.

26. Broer L, Codd V, Nyholt DR, Deelen J, Mangino M, Willemsen G, et al. Meta-analysis of telomere length in 19,713 subjects reveals high heritability, stronger maternal inheritance and a paternal age effect. European journal of human genetics : EJHG 21:1163–8 (2013). doi:10.1038/ejhg.2012.303 PMID 23321625.

27. Codd V, Wang Q, Allara E, Musicha C, Kaptoge S, Stoma S, et al. Polygenic basis and biomedical consequences of telomere length variation. Nature genetics 53:1425–1433 (2021). doi:10.1038/s41588-021-00944-6 PMID 34611362.

28. Telomeres Mendelian Randomization Collaboration, Haycock PC, Burgess S, Nounu A, Zheng J, Okoli GN et al. Association Between Telomere Length and Risk of Cancer and Non-Neoplastic Diseases: A Mendelian Randomization Study. JAMA oncology 3:636-651 (2017). doi:10.1001/jamaoncol.2016.5945 PMID 28241208.

29. Demanelis K, Jasmine F, Chen LS, Chernoff M, Tong L, Delgado D, et al. Determinants of telomere length across human tissues. Science (New York, N.Y.) 369 (2020). doi:10.1126/science.aaz6876 PMID 32913074.

30. Aviv A, Shay JW Reflections on telomere dynamics and ageing-related diseases in humans. Philosophical transactions of the Royal Society of London. Series B, Biological sciences 373 (2018). doi:10.1098/rstb.2016.0436 PMID 29335375.

31. Dutta S Unveiling the telomere-p53-PGC ageing axis. Nature reviews. Endocrinology 19:686 (2023). doi:10.1038/s41574-023-00905-5 PMID 37773274.

32. Aran D, Sirota M, Butte AJ Systematic pan-cancer analysis of tumour purity. Nature communications 6:8971 (2015). doi:10.1038/ncomms9971 PMID 26634437.

33. Yoshihara K, Shahmoradgoli M, Martínez E, Vegesna R, Kim H, Torres-Garcia W, et al. Inferring tumour purity and stromal and immune cell admixture from expression data. Nature communications 4:2612 (2013). doi:10.1038/ncomms3612 PMID 24113773.

34. Thorsson V, Gibbs DL, Brown SD, Wolf D, Bortone DS, Ou Yang TH, et al. The Immune Landscape of Cancer. Immunity 48:812–830.e14 (2018). doi:10.1016/j.immuni.2018.03.023 PMID 29628290.

35. Newman AM, Liu CL, Green MR, Gentles AJ, Feng W, Xu Y, et al. Robust enumeration of cell subsets from tissue expression profiles. Nature methods 12:453–7 (2015). doi:10.1038/nmeth.3337 PMID 25822800.

36. Livingstone J, Shiah YJ, Yamaguchi TN, Heisler LE, Huang V, Lesurf R, et al. The telomere length landscape of prostate cancer. Nature communications 12:6893 (2021). doi:10.1038/s41467-021-27223-6 PMID 34824250.

37. Hu K, Ghandi M, Huang FW Integrated evaluation of telomerase activation and telomere maintenance across cancer cell lines. eLife 10 (2021). doi:10.7554/eLife.66198 PMID 34486523.

38. Carter SL, Cibulskis K, Helman E, McKenna A, Shen H, Zack T, et al. Absolute quantification of somatic DNA alterations in human cancer. Nature biotechnology 30:413–21 (2012). doi:10.1038/nbt.2203 PMID 22544022.

39. Benjamini Y, Hochberg Y Controlling the False Discovery Rate: A Practical and Powerful Approach to Multiple Testing. Journal of the Royal Statistical Society Series B: Statistical Methodology 57:289–300 (1995). doi:10.1111/j.2517-6161.1995.tb02031.x.

40. Racle J, de Jonge K, Baumgaertner P, Speiser DE, Gfeller D Simultaneous enumeration of cancer and immune cell types from bulk tumor gene expression data. eLife 6 (2017). doi:10.7554/eLife.26476 PMID 29130882.

41. Mundlak Y On the Pooling of Time Series and Cross Section Data. Econometrica 46:69 (1978). doi:10.2307/1913646.

42. Carrot-Zhang J, Chambwe N, Damrauer JS, Knijnenburg TA, Robertson AG, Yau C, et al. Comprehensive Analysis of Genetic Ancestry and Its Molecular Correlates in Cancer. Cancer cell 37:639–654.e6 (2020). doi:10.1016/j.ccell.2020.04.012 PMID 32396860.

43. Ceccarelli M, Barthel FP, Malta TM, Sabedot TS, Salama SR, Murray BA, et al. Molecular Profiling Reveals Biologically Discrete Subsets and Pathways of Progression in Diffuse Glioma. Cell 164:550–63 (2016). doi:10.1016/j.cell.2015.12.028 PMID 26824661.

44. Cancer Genome Atlas Research Network, Brat DJ, Verhaak RG, Aldape KD, Yung WK, Salama SR, et al. Comprehensive, Integrative Genomic Analysis of Diffuse Lower-Grade Gliomas. The New England journal of medicine 372:2481-98 (2015). doi:10.1056/NEJMoa1402121 PMID 26061751.

45. Louis DN, Perry A, Wesseling P, Brat DJ, Cree IA, Figarella-Branger D, et al. The 2021 WHO Classification of Tumors of the Central Nervous System: a summary. Neuro-oncology 23:1231–1251 (2021). doi:10.1093/neuonc/noab106 PMID 34185076.

46. Barthel FP, Johnson KC, Varn FS, Moskalik AD, Tanner G, Kocakavuk E, et al. Longitudinal molecular trajectories of diffuse glioma in adults. Nature 576:112–120 (2019). doi:10.1038/s41586-019-1775-1 PMID 31748746.

47. Liu J, Lichtenberg T, Hoadley KA, Poisson LM, Lazar AJ, Cherniack AD, et al. An Integrated TCGA Pan-Cancer Clinical Data Resource to Drive High-Quality Survival Outcome Analytics. Cell 173:400–416.e11 (2018). doi:10.1016/j.cell.2018.02.052 PMID 29625055.

48. Hoadley KA, Yau C, Hinoue T, Wolf DM, Lazar AJ, Drill E, et al. Cell-of-Origin Patterns Dominate the Molecular Classification of 10,000 Tumors from 33 Types of Cancer. Cell 173:291–304.e6 (2018). doi:10.1016/j.cell.2018.03.022 PMID 29625048.

49. Wang Q, Zhan Y, Pedersen NL, Fang F, Hägg S Telomere Length and All-Cause Mortality: A Meta-analysis. Ageing research reviews 48:11–20 (2018). doi:10.1016/j.arr.2018.09.002 PMID 30254001.

50. Xiong F, Frasch WD ΩqPCR measures telomere length from single-cells in base pair units. Nucleic acids research 49:e120 (2021). doi:10.1093/nar/gkab753 PMID 34534325.

51. Meeker AK, Gage WR, Hicks JL, Simon I, Coffman JR, Platz EA, et al. Telomere length assessment in human archival tissues: combined telomere fluorescence in situ hybridization and immunostaining. The American journal of pathology 160:1259–68 (2002). doi:10.1016/S0002-9440(10)62553-9 PMID 11943711.

52. Dutta S, Goli A Beyond static biomarkers: systems biology and AI for decoding cancer dynamics. Frontiers in systems biology 6:1855016 (2026). doi:10.3389/fsysb.2026.1855016 PMID 42523586.

53. Cerami E, Gao J, Dogrusoz U, Gross BE, Sumer SO, Aksoy BA, et al. The cBio cancer genomics portal: an open platform for exploring multidimensional cancer genomics data. Cancer discovery 2:401–4 (2012). doi:10.1158/2159-8290.CD-12-0095 PMID 22588877.

54. Gao J, Aksoy BA, Dogrusoz U, Dresdner G, Gross B, Sumer SO, et al. Integrative analysis of complex cancer genomics and clinical profiles using the cBioPortal. Science signaling 6:pl1 (2013). doi:10.1126/scisignal.2004088 PMID 23550210.

55. Harris CR, Millman KJ, van der Walt SJ, Gommers R, Virtanen P, Cournapeau D, et al. Array programming with NumPy. Nature 585:357–362 (2020). doi:10.1038/s41586-020-2649-2 PMID 32939066.

56. Virtanen P, Gommers R, Oliphant TE, Haberland M, Reddy T, Cournapeau D, et al. SciPy 1.0: fundamental algorithms for scientific computing in Python. Nature methods 17:261–272 (2020). doi:10.1038/s41592-019-0686-2 PMID 32015543.

57. Hunter J Matplotlib: A 2D Graphics Environment. Computing in Science & Engineering 9:90-95 (2007). doi:10.1109/MCSE.2007.55.

