## Supplementary File for "A replicated patient-specific component of tumour telomere length across two pan-cancer cohorts"

**Subhajit Dutta**

Department of Biochemistry and Molecular Cell Biology (IBMZ), Center for Experimental Medicine, University Medical Center Hamburg-Eppendorf, Hamburg, Germany

#### Contents

**Supplementary Figures**


- Supplementary Figure S1. Sample selection, barcode harmonisation and merge quality control
- Supplementary Figure S2. Purity-estimator sensitivity, conditioning, and the circularity between consensus purity and leukocyte fraction
- Supplementary Figure S3. The age/*IDH* glioma analysis, shown in full as a worked negative
- Supplementary Figure S4. Two arms that did not support a claim: cell-line cross-validation and clinical outcome
- Supplementary Figure S5. Why the between-cohort coefficient is not an efficiency of transmission, and the limits of a germline test at this sample size


**Supplementary Tables**


- Table S1. Attrition from the published resource to the primary analysis set
- Table S2. Cohort sizes and the inclusion threshold
- Table S3. Per-cohort matched-normal correlation, TCGA
- Table S4. Per-cohort matched-normal correlation, PCAWG
- Table S5. Two-level within/between decomposition
- Table S6. Variance of tumour telomere length partitioned by level
- Table S7. Telomere attrition with age by sample type and assay
- Table S7b. Robustness of the age calibration to cohort, centre and sex
- Table S8. Per-cohort correlation of telomere length with tumour purity
- Table S9. Immune-subset associations with telomere length
- Table S10. Published telomere associations before and after adjustment
- Table S11. Purity estimators: circularity with leukocyte fraction and its consequence
- Table S12. Normal→tumour coefficient by purity quartile — the admixture falsification test
- Table S13. Leave-one-cohort-out stability of the between-cohort coefficient
- Table S14. Discrimination of telomere length for progression-free interval
- Table S15. Instability of CCLE lineage medians
- Table S16. Decomposition refitted by matched-normal type
- Table S17. Genetic ancestry as a germline proxy, and the power available
- Table S18. The association with technical structure entered as fixed effects
- Table S19. Nested demographic and compositional sensitivity, including sex


**Supplementary Notes**


- Note 1. Barcode harmonisation and the merge procedure
- Note 2. Why the analysis set is restricted to whole-genome and low-pass data
- Note 3. Circularity among tumour-purity estimators
- Note 4. Verification procedures: figure auditing and citation checking


References follow the numbering of the main text.

### Supplementary Figures

#### Supplementary Figure S1

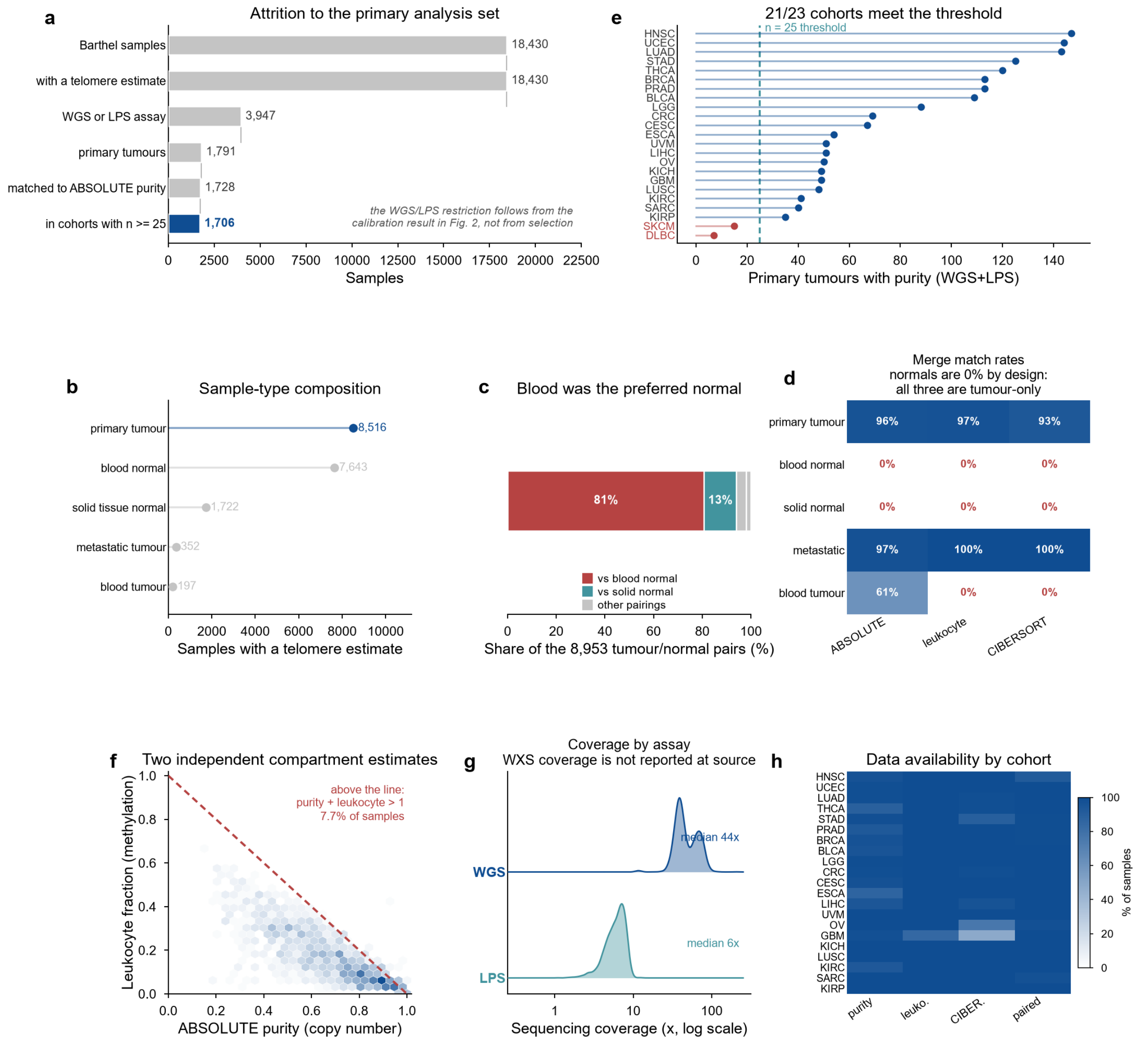


**Supplementary Figure S1. Sample selection, barcode harmonisation and merge quality control.**

(**a**) Attrition from the 18,430 samples of the published resource to the 1,706 primary tumours analysed here. The restriction to whole-genome and low-pass libraries follows from the age calibration in Fig. 2 and is not a selection made for convenience. (**b**) Sample-type composition of the resource, by TCGA sample-type code. (**c**) Which normal tissue was used as the matched reference across the 8,953 published tumour/normal pairs; blood was preferred wherever both were available. (**d**) Match rate of each external annotation to each sample type after barcode harmonisation. Normals match at 0% because ABSOLUTE, leukocyte fraction and CIBERSORT are all defined for tumour samples only — this is by design, not merge failure. (**e**) Primary tumours with a purity call per cohort; the dashed line is the n = 25 inclusion threshold and the two excluded cohorts are shown in red. (**f**) The two compartment estimates used in this work, ABSOLUTE purity (copy number) and leukocyte fraction (methylation), are methodologically independent. Points above the dashed line sum to more than one and are therefore mutually inconsistent; they are 7.7% of samples and are retained rather than filtered. (**g**) Sequencing coverage by assay. Coverage is not reported for exome samples at source, which is why the coverage covariate exists only for the whole-genome and low-pass set. (**h**) Availability of each annotation by cohort.

#### Supplementary Figure S2

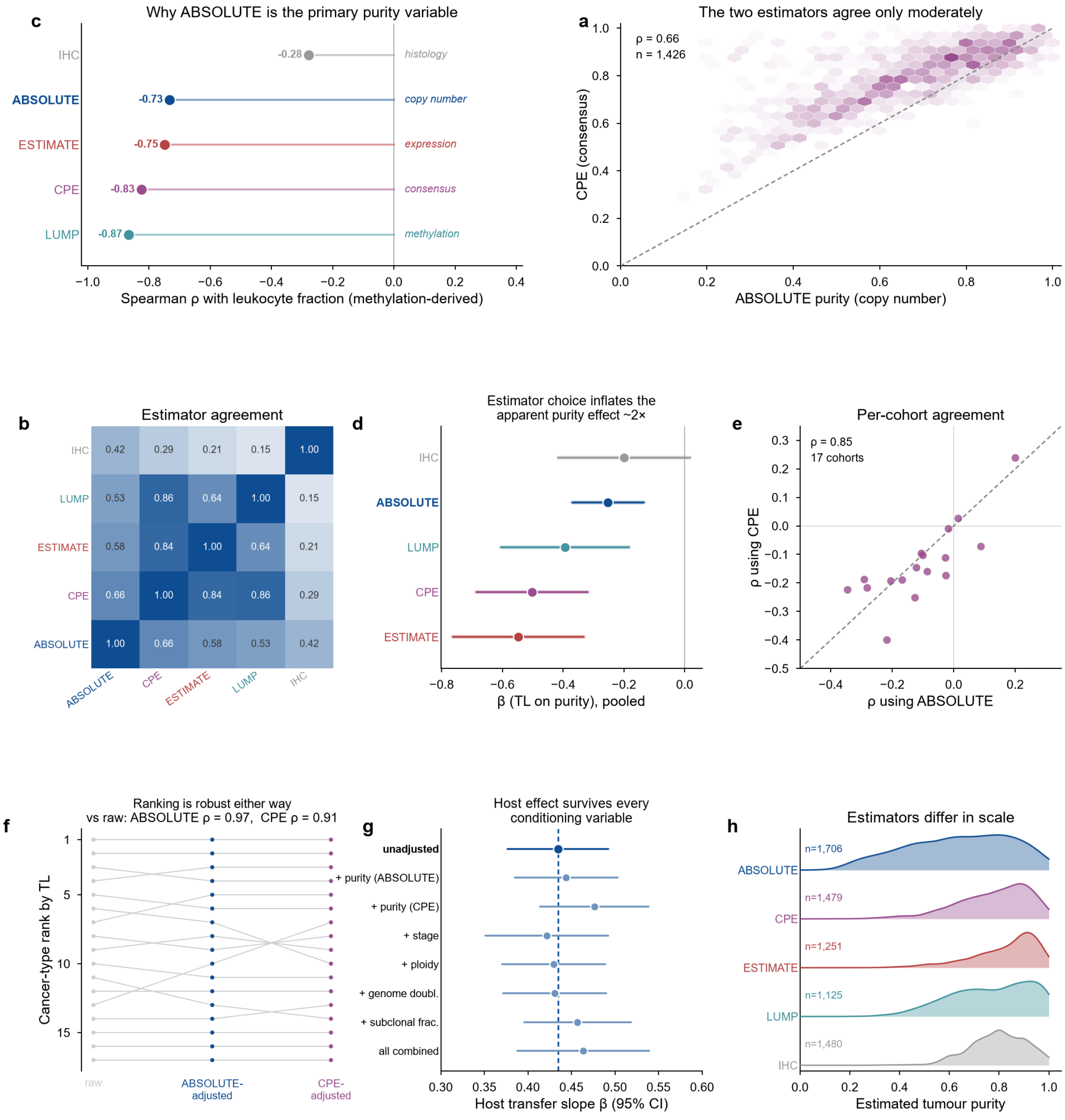


**Supplementary Figure S2. Purity-estimator sensitivity, conditioning, and the circularity between consensus purity and leukocyte fraction.**

(**a**) ABSOLUTE purity against the Consensus Purity Estimate for the samples carrying both; the dashed line is the identity. (**b**) Pairwise Spearman agreement among the five purity estimators. (**c**) Correlation of each purity estimator with methylation-derived leukocyte fraction, annotated with the measurement modality each estimator uses. LUMP is itself derived from leukocyte-specific unmethylation and CPE contains LUMP as a component, so their strong negative correlations are partly definitional rather than biological. (**d**) Pooled effect of purity on telomere length under each estimator. The ordering follows the degree of shared measurement input in **c**, which is what circularity predicts. (**e**) Per-cohort effect estimated with ABSOLUTE against the same estimated with CPE. (**f**) Cross-cancer ranking, raw and after adjustment under each estimator. (**g**) The normal→tumour coefficient under nested conditioning; the dashed line is the unadjusted estimate. (**h**) Marginal distribution of each estimator, showing that they differ in scale as well as in correlation.

#### Supplementary Figure S3

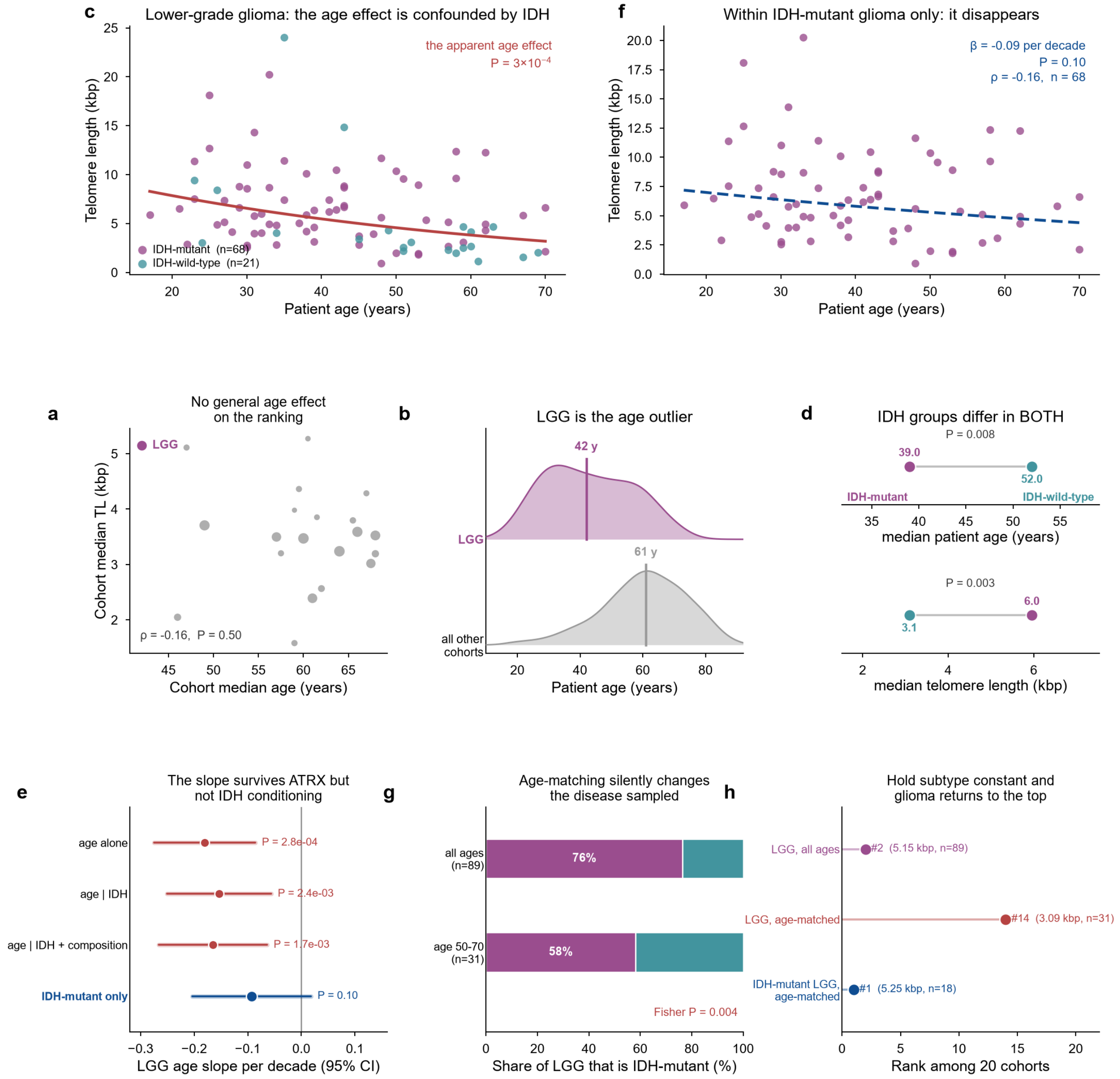

**Supplementary Figure S3. The age/*IDH* glioma analysis, shown in full as a worked negative.**

(**a**) Cohort median patient age against cohort median telomere length; there is no general age effect on the cross-cancer ranking. (**b**) Age distribution of lower-grade glioma against all other cohorts pooled; LGG is a marked outlier. (**c**) Telomere length against age within lower-grade glioma, with *IDH* status revealed by colour. The apparent age effect is significant, but the two *IDH* groups occupy different regions of the age axis. (**d**) *IDH*-mutant and *IDH*-wild-type gliomas differ significantly in patient age **and** in telomere length, which is the definition of a confounder for this comparison. (**e**) The LGG age slope under nested conditioning; it survives adjustment for composition but not for *IDH* status. (**f**) The decisive control: within *IDH*-mutant gliomas alone the age slope is not significant. (**g**) Restricting to ages 50–70 silently changes the molecular composition of the cohort, lowering the *IDH*-mutant share. The age-matched cohort is therefore a different disease, not the same disease at a different age. (**h**) Consequence for the ranking: holding molecular subtype constant returns glioma to the top.

#### Supplementary Figure S4

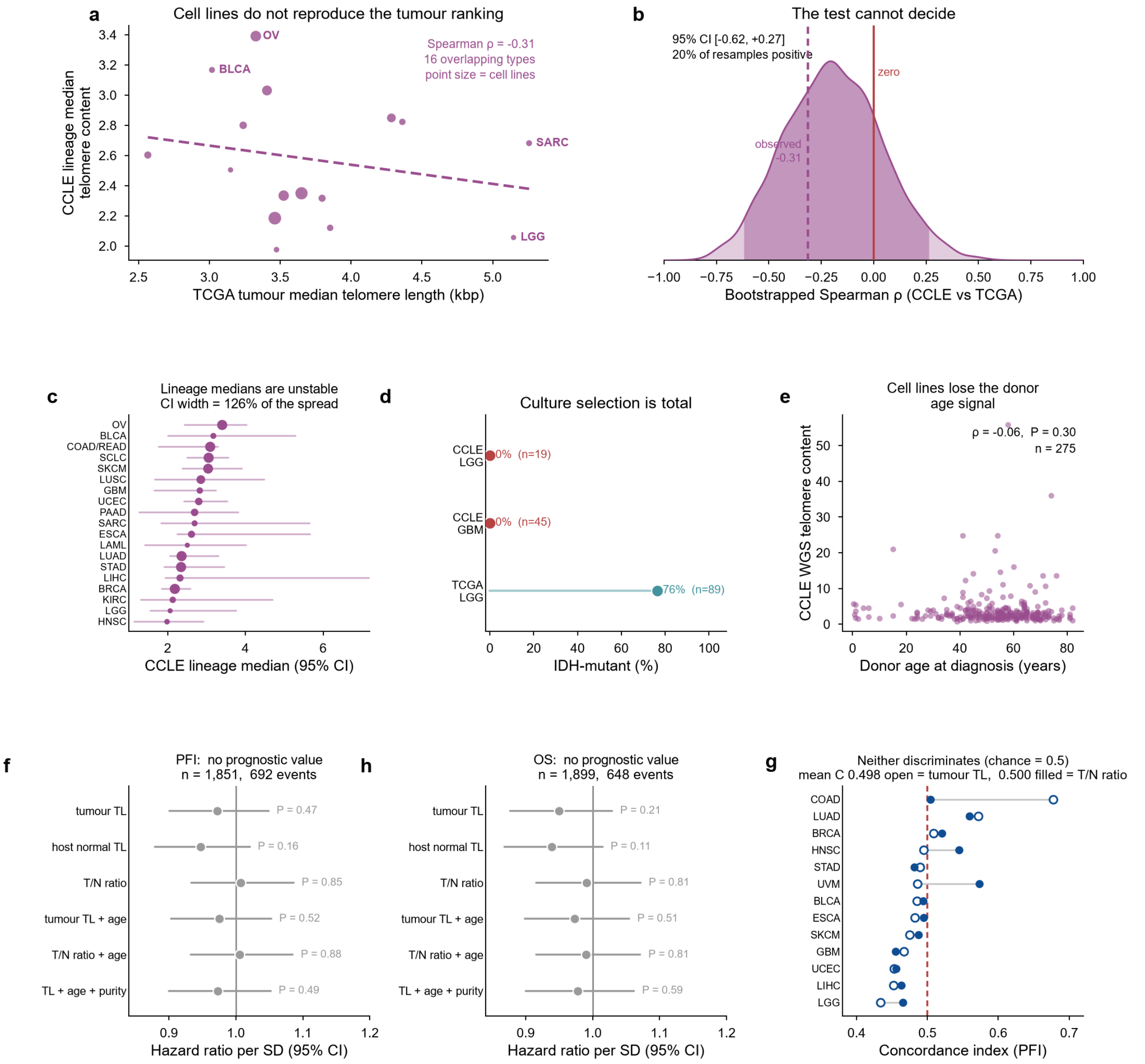


**Supplementary Figure S4. Two arms that did not support a claim: cell-line cross-validation and clinical outcome.**

(**a**) CCLE lineage median telomere content against TCGA cancer-type median telomere length for the overlapping types. (**b**) Bootstrap distribution of that correlation. The interval spans zero and 20% of resamples are positive, so the comparison cannot distinguish the hypotheses in either direction; it is uninformative rather than negative. (**c**) Bootstrap confidence intervals on each CCLE lineage median. The median interval width is 126% of the entire between-lineage spread, and 15 of 19 lineages have an interval wider than that spread - the structural reason **b** cannot resolve. (**d**) *IDH1* mutation frequency in CCLE glioma lines against TCGA lower-grade glioma; culture selection is complete, so the cell-line panel does not contain the biology that drives the tumour result. (**e**) Donor age at diagnosis against cell-line telomere content — the host age signal is absent, consistent with immortalisation erasing it. (**f**, **h**) Cox hazard ratios per standard deviation for progression-free interval and overall survival, stratified by cancer type. (**g**) Per-cancer-type concordance index for tumour telomere length and for the tumour/normal ratio; the dashed line is chance.

#### Supplementary Figure S5

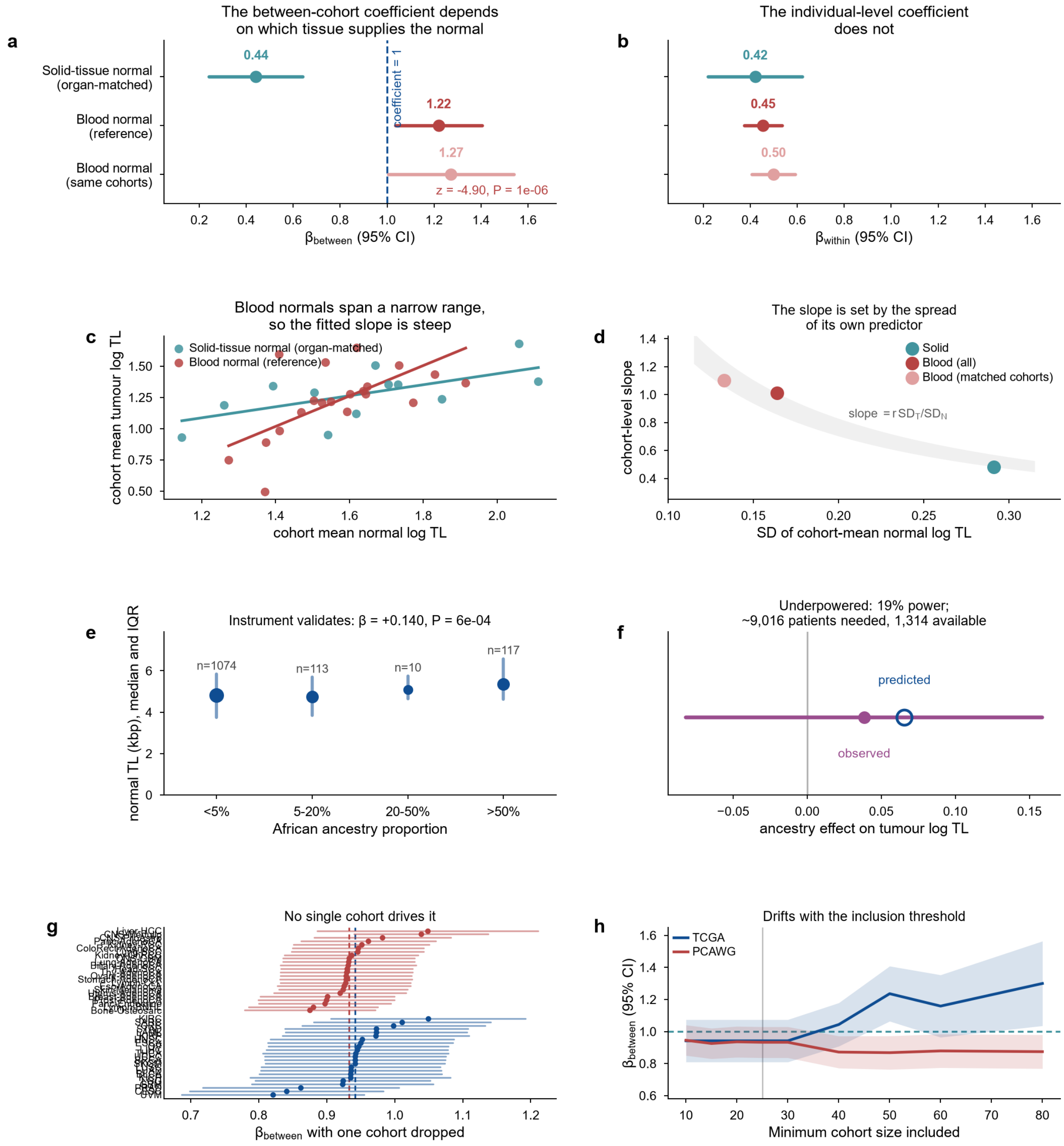


**Supplementary Figure S5. Why the between-cohort coefficient is not an efficiency of transmission, the limits of a germline test at this sample size, and the diagnostics for that coefficient.**

(**a**) The between-cohort coefficient refitted by matched-normal type. With an organ-matched solid-tissue reference it is 0.44 and excludes a coefficient of one; with a blood-normal reference it is 1.22. The third row restricts the blood arm to the solid arm's own cohorts, so the gap is not a difference in which cancers are included. (**b**) The same three arms for the individual-level coefficient, on the same scale: it does not move. (**c**) Cohort means with the fitted between-cohort slopes. Cohort-mean blood telomere length spans a narrow range, so the fitted slope is steep; cohort-mean tissue telomere length spans a wide one, so it is shallow. (**d**) The cohort-level regression slope against the spread of its own predictor. A least-squares slope equals $r$ × SD(tumour) / SD(normal) identically, so it must fall inside the shaded band, which spans the range of $r$ × SD(tumour) observed across the three arms. The size-weighted between-cohort coefficient of panel **a** tracks this quantity but is not identical to it. (**e**) Validity check for the germline analysis: African ancestry proportion against the patient's own normal telomere length, median and interquartile range, marker area scaled by the number of patients. The established association is recovered. (**f**) The tumour-side test itself: the observed ancestry effect on tumour telomere length with its 95% interval, against the effect the constitutional model predicts. The interval contains the prediction and zero alike, because the study has 19% power to separate them. (**g**) Leave-one-cohort-out estimates of the between-cohort coefficient, with the full-data value dashed. (**h**) Sensitivity of the between-cohort coefficient to the minimum cohort size. Panels **g** and **h** are influence and threshold diagnostics for a coefficient that panels **a**–**d** show is not an efficiency of transmission; they are reported for completeness.

#### Supplementary Tables

##### Table S1. Attrition from the published resource to the primary analysis set

| Selection step | Samples remaining | Change |
|---|---|---|
| Barthel samples | 18,430 | — |
| with a telomere estimate | 18,430 | 0 |
| WGS or LPS assay | 3,947 | −14,483 |
| primary tumours | 1,791 | −2,156 |
| matched to ABSOLUTE purity | 1,728 | −63 |
| in cohorts with n ≥ 25 | 1,706 | −22 |

The restriction to whole-genome and low-pass libraries is a consequence of the age calibration reported in the main text, not a convenience filter; the exome samples fail a biological sanity check rather than a quality threshold.

##### Table S2. Cohort sizes and the inclusion threshold

| TCGA cohort | Primary tumours with purity (WGS+LPS) | Status |
|---|---|---|
| HNSC | 147 | included |
| UCEC | 144 | included |
| LUAD | 143 | included |
| STAD | 125 | included |
| THCA | 120 | included |
| PRAD | 113 | included |
| BRCA | 113 | included |
| BLCA | 109 | included |
| LGG | 88 | included |
| CRC | 69 | included |
| CESC | 67 | included |
| ESCA | 54 | included |
| UVM | 51 | included |
| LIHC | 51 | included |
| OV | 50 | included |
| GBM | 49 | included |
| KICH | 49 | included |
| LUSC | 48 | included |
| KIRC | 41 | included |
| SARC | 40 | included |
| KIRP | 35 | included |
| SKCM | 15 | **excluded** (n < 25) |
| DLBC | 7 | **excluded** (n < 25) |

##### Table S3. Per-cohort matched-normal correlation, TCGA

| TCGA cohort | n | Spearman ρ | 95% CI | P |
|---|---|---|---|---|
| PRAD | 121 | 0.597 | 0.468 to 0.702 | $5.1 \times 10^{-13}$ |
| THCA | 135 | 0.499 | 0.351 to 0.627 | $7.3 \times 10^{-10}$ |
| LGG | 89 | 0.483 | 0.316 to 0.613 | $1.7 \times 10^{-6}$ |
| KIRP | 35 | 0.466 | 0.120 to 0.724 | 0.005 |
| UVM | 51 | 0.446 | 0.187 to 0.640 | 0.001 |
| HNSC | 139 | 0.424 | 0.273 to 0.556 | $2.0 \times 10^{-7}$ |

| TCGA cohort | n | Spearman ρ | 95% CI | P |
|---|---|---|---|---|
| KICH | 49 | 0.423 | 0.132 to 0.675 | 0.002 |
| LUAD | 144 | 0.345 | 0.197 to 0.482 | $2.3 \times 10^{-5}$ |
| BLCA | 114 | 0.319 | 0.150 to 0.477 | $5.4 \times 10^{-4}$ |
| LIHC | 54 | 0.297 | 0.008 to 0.541 | 0.029 |
| SKCM | 135 | 0.295 | 0.132 to 0.439 | $5.2 \times 10^{-4}$ |
| CESC | 69 | 0.279 | 0.042 to 0.498 | 0.020 |
| CRC | 70 | 0.266 | 0.041 to 0.469 | 0.026 |
| STAD | 126 | 0.258 | 0.067 to 0.432 | 0.004 |
| LUSC | 48 | 0.240 | −0.085 to 0.500 | 0.100 |
| ESCA | 62 | 0.226 | −0.033 to 0.482 | 0.077 |
| KIRC | 44 | 0.226 | −0.115 to 0.532 | 0.140 |
| SARC | 39 | 0.183 | −0.125 to 0.451 | 0.265 |
| BRCA | 116 | 0.163 | −0.034 to 0.341 | 0.080 |
| OV | 49 | 0.153 | −0.116 to 0.411 | 0.293 |
| GBM | 50 | 0.121 | −0.164 to 0.399 | 0.401 |
| UCEC | 146 | 0.114 | −0.047 to 0.270 | 0.171 |
| LAML | 50 | −0.012 | −0.310 to 0.309 | 0.936 |

22 of 23 cohorts show a positive matched-normal correlation. Confidence intervals are percentile bootstrap (2,000-5,000 resamples).

#### Table S4. Per-cohort matched-normal correlation, PCAWG

| PCAWG histology | n | Spearman ρ | 95% CI | P |
|---|---|---|---|---|
| Stomach-AdenoCA | 68 | 0.787 | 0.675 to 0.854 | $1.7 \times 10^{-15}$ |
| Biliary-AdenoCA | 33 | 0.619 | 0.283 to 0.845 | $1.2 \times 10^{-4}$ |
| Liver-HCC | 305 | 0.549 | 0.468 to 0.626 | $1.9 \times 10^{-25}$ |
| Breast-AdenoCA | 195 | 0.524 | 0.412 to 0.625 | $3.8 \times 10^{-15}$ |
| CNS-PiloAstro | 89 | 0.507 | 0.314 to 0.665 | $3.9 \times 10^{-7}$ |
| Head-SCC | 56 | 0.427 | 0.165 to 0.650 | 0.001 |
| Prost-AdenoCA | 178 | 0.422 | 0.287 to 0.536 | $4.6 \times 10^{-9}$ |
| Lymph-CLL | 90 | 0.401 | 0.201 to 0.579 | $9.0 \times 10^{-5}$ |
| Panc-AdenoCA | 229 | 0.384 | 0.263 to 0.498 | $1.8 \times 10^{-9}$ |
| Kidney-RCC | 143 | 0.375 | 0.218 to 0.513 | $3.9 \times 10^{-6}$ |
| Ovary-AdenoCA | 110 | 0.360 | 0.180 to 0.525 | $1.1 \times 10^{-4}$ |
| Thy-AdenoCA | 48 | 0.358 | 0.091 to 0.589 | 0.013 |
| Kidney-ChRCC | 43 | 0.342 | 0.025 to 0.624 | 0.025 |
| Lymph-BNHL | 107 | 0.300 | 0.104 to 0.478 | 0.002 |
| Eso-AdenoCA | 97 | 0.286 | 0.092 to 0.470 | 0.004 |
| Lung-AdenoCA | 37 | 0.282 | −0.061 to 0.568 | 0.090 |
| CNS-Medullo | 141 | 0.260 | 0.096 to 0.405 | 0.002 |
| Lung-SCC | 47 | 0.194 | −0.136 to 0.466 | 0.192 |
| Panc-Endocrine | 81 | 0.187 | −0.032 to 0.391 | 0.095 |
| CNS-GBM | 39 | 0.182 | −0.154 to 0.473 | 0.269 |
| Uterus-AdenoCA | 44 | 0.169 | −0.166 to 0.468 | 0.273 |
| Skin-Melanoma | 107 | 0.138 | −0.073 to 0.322 | 0.155 |
| ColoRect-AdenoCA | 52 | 0.107 | −0.158 to 0.365 | 0.450 |
| Bone-Osteosarc | 35 | 0.080 | −0.269 to 0.435 | 0.647 |

24 of 24 cohorts show a positive matched-normal correlation. Confidence intervals are percentile bootstrap (2,000-5,000 resamples).

##### Table S5. Two-level within/between decomposition

| Cohort | n | Groups | $\beta_{within}$ | $P_{within}$ | $\beta_{between}$ | $P_{between}$ | P ($\beta_w = \beta_b$) | P ($\beta_b = 1$) |
|---|---|---|---|---|---|---|---|---|
| TCGA (TelSeq, Barthel 2017) | 1,935 | 23 | 0.417 | $8.2 \times 10^{-31}$ | 0.942 | $2.2 \times 10^{-42}$ | $7.8 \times 10^{-12}$ | 0.388 |
| PCAWG (TelomereHunter, Sieverling 2020) | 2,374 | 24 | 0.671 | $5.8 \times 10^{-102}$ | 0.933 | $9.9 \times 10^{-75}$ | $5.6 \times 10^{-6}$ | 0.171 |

$\beta_{within}$ is the coefficient on the patient's deviation from their cohort mean; $\beta_{between}$ is the coefficient on the cohort mean itself. $\beta_{between}$ is a statistical between-cohort term, **not** an identified parameter of tissue transmission: as shown in Supplementary Fig. S5 and Table S16, its magnitude depends on which tissue supplies the matched normal and on the spread of that predictor. The final column tests it against a slope of one, which is a reference value and not a biological target.

##### Table S6. Variance of tumour telomere length partitioned by level

| Cohort | n | Within-cohort (%) | Between-cohort (%) | Residual (%) | Both terms (%) |
|---|---|---|---|---|---|
| TCGA | 1,935 | 6.1 | 8.6 | 85.3 | 14.7 |
| PCAWG | 2,374 | 15.7 | 11.1 | 73.2 | 26.8 |

The between-cohort share is reported for completeness. As shown in Supplementary Fig. S5 and Table S16, the between-cohort coefficient is not identified as a tissue-of-origin contribution, so this column should not be read as the contribution of the tissue of origin. The within-cohort share is the interpretable one.

##### Table S7. Telomere attrition with age by sample type and assay

| Sample type | Assay | n | bp per year | 95% CI | P |
|---|---|---|---|---|---|
| Blood normal | WGS | 730 | −30.05 | −38.3 to −21.8 | $1.9 \times 10^{-12}$ |
| Blood normal | LPS | 827 | −22.30 | −29.9 to −14.7 | $1.4 \times 10^{-8}$ |
| Blood normal | WXS | 6,015 | 2.66 (wrong sign) | 1.1 to 4.2 | $6.0 \times 10^{-4}$ |
| Solid tissue normal | WGS | 272 | −37.77 | −52.3 to −23.2 | $5.8 \times 10^{-7}$ |
| Solid tissue normal | LPS | 146 | −35.97 | −58.0 to −14.0 | 0.002 |
| Solid tissue normal | WXS | 1,300 | 8.04 (wrong sign) | 2.4 to 13.7 | 0.005 |
| Primary tumour | WGS | 924 | −14.72 | −35.0 to 5.6 | 0.156 |
| Primary tumour | LPS | 857 | −16.07 | −30.4 to −1.7 | 0.028 |
| Primary tumour | WXS | 6,668 | 2.18 (wrong sign) | 0.6 to 3.8 | 0.008 |

Published cross-sectional leukocyte telomere attrition in adults is approximately −20 to −30 bp per year [24,25]. Whole-genome and low-pass estimates fall inside that range in blood normals; the unadjusted exome estimates do not, and Table S7b shows what survives adjustment.

##### Table S7b. Robustness of the age calibration to cohort, centre and sex

| Analysis | Assay | Model | n | bp per year | 95% CI | P |
|---|---|---|---|---|---|---|

| Analysis | Assay | Model | n | bp per year | 95% CI | P |
|---|---|---|---|---|---|---|
| Marginal / covariate-adjusted | WGS | unadjusted | 730 | −30.05 | −38.3 to −21.8 | $1.9 \times 10^{-12}$ |
| Marginal / covariate-adjusted | WGS | + cancer type, centre, sex | 730 | −29.43 | −37.7 to −21.2 | $5.7 \times 10^{-12}$ |
| Marginal / covariate-adjusted | LPS | unadjusted | 827 | −22.30 | −29.9 to −14.7 | $1.4 \times 10^{-8}$ |
| Marginal / covariate-adjusted | LPS | + cancer type, centre, sex | 827 | −28.91 | −37.1 to −20.7 | $8.2 \times 10^{-12}$ |
| Marginal / covariate-adjusted | WXS | unadjusted | 6,015 | 2.66 | 1.1 to 4.2 | $6.0 \times 10^{-4}$ |
| Marginal / covariate-adjusted | WXS | + cancer type, centre, sex | 6,015 | −0.60 | −2.1 to 0.9 | 0.448 |
| Within-cancer meta-analysis | WGS | 14 cohorts, inverse-variance | 620 | −33.12 | −40.8 to −25.5 | $< 1 \times 10^{-300}$ |
| Within-cancer meta-analysis | LPS | 12 cohorts, inverse-variance | 810 | −27.33 | −34.7 to −20.0 | $3.7 \times 10^{-13}$ |
| Within-cancer meta-analysis | WXS | 29 cohorts, inverse-variance | 6,015 | −0.14 | −0.3 to $3.4 \times 10^{-3}$ | 0.055 |
| Same-aliquot, both assays | WGS | paired WGS vs WXS aliquots | 684 | −38.94 | −47.8 to −30.0 | $5.4 \times 10^{-17}$ |
| Same-aliquot, both assays | WXS | paired WGS vs WXS aliquots | 684 | 2.58 | −6.7 to 11.9 | 0.585 |
| Same-aliquot, both assays | WGS-WXS difference | paired difference on age | 684 | −41.52 | −54.3 to −28.7 | $3.3 \times 10^{-10}$ |

Assay usage, patient age and cancer type are not randomly distributed across TCGA, so a marginal slope could in principle be produced by between-cohort composition rather than by the assay. Three tests of increasing stringency exclude that. Adjusting for cancer type, sequencing centre and sex leaves the whole-genome and low-pass slopes essentially unchanged and moves the exome slope to −0.6 bp per year, indistinguishable from zero. Estimating the slope inside each cancer type and pooling by inverse-variance meta-analysis — where between-cohort composition cannot contribute at all — gives the same picture. The decisive comparison is the last: for 684 blood-normal aliquots the identical DNA was sequenced by both assays, so patient, age, cancer type, centre and material are constant and only the assay differs; the whole-genome estimate declines at −38.9 bp per year and the exome estimate from the same DNA does not decline. The positive sign in the unadjusted exome slope is therefore not robust, but the absence of any age signal is, and it is the more consequential defect. Low-pass blood normals come from a single sequencing centre, so no centre term is estimable within that assay.

#### Table S8. Per-cohort correlation of telomere length with tumour purity

| Cohort | n | Spearman ρ (TL vs purity) | 95% CI | P | q (BH) |
|---|---|---|---|---|---|
| CRC | 69 | −0.346 | −0.549 to −0.109 | 0.004 | 0.019 |
| BRCA | 113 | −0.292 | −0.463 to −0.106 | 0.002 | 0.018 |
| UCEC | 144 | −0.282 | −0.430 to −0.123 | $6.3 \times 10^{-4}$ | 0.013 |

| Cohort | n | Spearman ρ (TL vs purity) | 95% CI | P | q (BH) |
|---|---|---|---|---|---|
| STAD | 125 | −0.266 | −0.428 to −0.094 | 0.003 | 0.019 |
| KIRC | 41 | −0.219 | −0.518 to 0.120 | 0.170 | 0.356 |
| PRAD | 113 | −0.205 | −0.400 to 0.004 | 0.029 | 0.124 |
| ESCA | 54 | −0.195 | −0.447 to 0.076 | 0.157 | 0.356 |
| BLCA | 109 | −0.169 | −0.353 to 0.021 | 0.079 | 0.278 |
| CESC | 67 | −0.128 | −0.350 to 0.110 | 0.304 | 0.531 |
| LUAD | 143 | −0.121 | −0.286 to 0.042 | 0.149 | 0.356 |
| LIHC | 51 | −0.105 | −0.405 to 0.195 | 0.462 | 0.640 |
| OV | 50 | −0.101 | −0.354 to 0.168 | 0.487 | 0.640 |
| THCA | 120 | −0.086 | −0.273 to 0.102 | 0.350 | 0.565 |
| KIRP | 35 | −0.028 | −0.356 to 0.284 | 0.873 | 0.928 |
| HNSC | 147 | −0.026 | −0.198 to 0.147 | 0.751 | 0.928 |
| UVM | 51 | −0.021 | −0.309 to 0.282 | 0.883 | 0.928 |
| GBM | 49 | −0.018 | −0.292 to 0.265 | 0.904 | 0.928 |
| LUSC | 48 | 0.013 | −0.281 to 0.298 | 0.928 | 0.928 |
| LGG | 88 | 0.088 | −0.127 to 0.300 | 0.418 | 0.626 |
| KICH | 49 | 0.200 | −0.096 to 0.475 | 0.167 | 0.356 |
| SARC | 40 | 0.205 | −0.134 to 0.512 | 0.203 | 0.388 |

#### Table S9. Immune-subset associations with telomere length

| Immune subset | β per SD | SE | Cohorts | $I^2$ (%) | P | q (BH) |
|---|---|---|---|---|---|---|
| B cells naive | 0.0458 | 0.0116 | 18 | 0.0 | $7.5 \times 10^{-5}$ | 0.001 |
| T cells follicular helper | 0.0378 | 0.0112 | 20 | 42.2 | $7.4 \times 10^{-4}$ | 0.007 |
| T cells CD4 memory resting | 0.0356 | 0.0117 | 20 | 33.4 | 0.002 | 0.015 |
| B cells memory | 0.0301 | 0.0120 | 16 | 51.4 | 0.012 | 0.056 |
| Plasma cells | 0.0215 | 0.0111 | 19 | 21.2 | 0.053 | 0.168 |
| Mast cells resting | −0.0217 | 0.0109 | 19 | 0.0 | 0.047 | 0.168 |
| T cells CD8 | 0.0217 | 0.0117 | 20 | 48.4 | 0.063 | 0.172 |
| Neutrophils | 0.0213 | 0.0135 | 14 | 0.0 | 0.115 | 0.274 |
| T cells regulatory Tregs | 0.0168 | 0.0121 | 18 | 50.2 | 0.166 | 0.350 |
| Monocytes | 0.0132 | 0.0109 | 20 | 50.0 | 0.225 | 0.427 |
| Macrophages M2 | 0.0139 | 0.0122 | 20 | 36.7 | 0.256 | 0.442 |
| Macrophages M0 | −0.0119 | 0.0116 | 18 | 0.0 | 0.302 | 0.446 |
| Dendritic cells activated | −0.0116 | 0.0113 | 17 | 0.0 | 0.305 | 0.446 |
| NK cells activated | −0.0079 | 0.0112 | 20 | 0.0 | 0.481 | 0.652 |
| Dendritic cells resting | −0.0058 | 0.0125 | 14 | 24.1 | 0.642 | 0.813 |
| T cells CD4 memory activated | 0.0018 | 0.0200 | 7 | 0.0 | 0.927 | 0.927 |
| Mast cells activated | 0.0043 | 0.0167 | 10 | 0.0 | 0.795 | 0.927 |
| NK cells resting | −0.0013 | 0.0118 | 16 | 12.3 | 0.914 | 0.927 |
| Macrophages M1 | 0.0012 | 0.0119 | 20 | 0.0 | 0.923 | 0.927 |

Absolute subset abundance (leukocyte fraction × CIBERSORT relative fraction), adjusted for tumour purity and patient age, meta-analysed across cohorts by inverse-variance weighting. Effects are per standard deviation of abundance, against a within-cohort telomere-length standard deviation of 57%.

##### Table S10. Published telomere associations before and after adjustment

| Association | n | β raw | q raw | β adjusted | q adjusted | Attenuation (%) | Verdict |
|---|---|---|---|---|---|---|---|
| ATRXDAXXstatus=ATRX/DAXX wt | 1,506 | −0.386 | $1.9 \times 10^{-12}$ | −0.368 | $2.0 \times 10^{-11}$ | 4.8 | HOLDS |
| TERTexprStatus=TERT expr | 1,506 | −0.256 | $1.5 \times 10^{-14}$ | −0.269 | $5.9 \times 10^{-15}$ | −4.9 | HOLDS |
| ATRXstatus=ATRX wt | 1,506 | −0.424 | $5.5 \times 10^{-14}$ | −0.407 | $6.3 \times 10^{-13}$ | 4.1 | HOLDS |
| TERTstatus=TERTp meth | 441 | −0.109 | 0.051 | −0.096 | 0.101 | — | null both |
| TERTexpr | 1,553 | 0.044 | 0.003 | 0.047 | 0.002 | 5.7 | HOLDS |
| ATRXexpr | 1,553 | −0.011 | 0.513 | −0.005 | 0.745 | — | null both |
| TelomeraseSignatureScore | 1,553 | 0.011 | 0.513 | −0.005 | 0.745 | — | null both |
| TERCexpr | 1,553 | $-6.7 \times 10^{-4}$ | 0.962 | 0.005 | 0.745 | — | null both |

Adjustment is for ABSOLUTE purity, methylation-derived leukocyte fraction and patient age. No association is gained or lost. Percentage attenuation is shown only for the four associations that hold; for coefficients indistinguishable from zero the ratio has a near-zero denominator and is not interpretable, so it is omitted.

##### Table S11. Purity estimators: circularity with leukocyte fraction and its consequence

| Purity estimator | Measurement modality | n | ρ with leukocyte fraction | Pooled β (TL on purity) | 95% CI | P |
|---|---|---|---|---|---|---|
| LUMP | methylation | 1,125 | −0.866 | −0.394 | −0.605 to −0.183 | $2.5 \times 10^{-4}$ |
| CPE | consensus | 1,470 | −0.825 | −0.502 | −0.685 to −0.319 | $7.7 \times 10^{-8}$ |
| ESTIMATE | expression | 1,246 | −0.748 | −0.548 | −0.764 to −0.333 | $5.9 \times 10^{-7}$ |
| ABSOLUTE | copy number | 1,697 | −0.733 | −0.253 | −0.370 to −0.135 | $2.6 \times 10^{-5}$ |
| IHC | histology | 1,471 | −0.278 | −0.199 | −0.416 to 0.018 | 0.072 |

Estimators are ordered by their correlation with leukocyte fraction. The pooled purity effect on telomere length tracks that ordering, roughly doubling from the least to the most entangled estimator. ABSOLUTE is used as primary throughout because copy number is the modality least shared with the immune covariates.

##### Table S12. Normal→tumour coefficient by purity quartile — the admixture falsification test

| Purity quartile | n | Purity range | Median purity | Normal→tumour β (raw scale) | 95% CI | β under pure admixture |
|---|---|---|---|---|---|---|
| Q1 | 463 | 0.08–0.48 | 0.38 | 0.426 | 0.338 to 0.514 | 0.620 |
| Q2 | 468 | 0.49–0.66 | 0.58 | 0.351 | 0.196 to 0.506 | 0.420 |
| Q3 | 437 | 0.67–0.81 | 0.74 | 0.357 | 0.235 to 0.478 | 0.260 |
| Q4 | 456 | 0.82–1.00 | 0.90 | 0.360 | 0.048 to 0.672 | 0.100 |

All coefficients are raw-scale. Predicted β(Q4)/β(Q1) under pure admixture: **0.161**; observed: **0.845**. The quartile prediction uses median purities and is an intuitive approximation; the formal test is the continuous model below. Raw-scale interaction β = −0.092, P = 0.711.

**Formal test against the pure-admixture null.** Under Y = pT + (1−p)H with tumour and host telomere length independent, the conditional model predicts $\beta_H$ = 1 and β~H×p~ = −1, not merely a zero interaction. Observed: $\beta_H$ = 0.435 (SE 0.170, P = $9.4 \times 10^{-4}$ against 1) and β~H×p~ = −0.092 (SE 0.248, P = $2.6 \times 10^{-4}$ against −1). Joint Wald test: F(2, 1,820) = 6.87, **P = 0.001**, n = 1,824. The linear main effect of purity assumes p·E[T | p] is adequately represented; repeating the joint test with a natural cubic spline in purity (4 df) gives F = 6.10, P = 0.002, and with a quadratic F = 6.10, P = 0.002, so the rejection does not depend on that assumption.

The admixture identity is a statement about raw telomere length, so the coefficient is fitted on the raw scale, where a pure two-compartment model requires it to equal (1 − purity) and therefore to fall by a factor of 0.16 across these quartiles. The observed ratio is 0.85, a fall of 15% against a required 84%, and the raw-scale interaction with purity is not significant. Elsewhere in this work telomere length is modelled on the log scale; there the correct expectation under pure admixture is (1 − purity) × $TL_{host}/TL_{measured}$, not (1 − purity), which is why the falsification is fitted on the raw scale here. Pure admixture is therefore excluded as the sole explanation; a partial admixture contribution alongside a patient-specific association is not.

#### Table S13. Leave-one-cohort-out stability of the between-cohort coefficient

| Cohort | Cohorts dropped (one at a time) | Minimum $\beta_{between}$ | Maximum $\beta_{between}$ | Full-data $\beta_{between}$ |
|---|---|---|---|---|
| TCGA | 23 | 0.821 | 1.049 | 0.942 |
| PCAWG | 24 | 0.875 | 1.049 | 0.933 |

#### Table S14. Discrimination of telomere length for progression-free interval

| Cancer type | n | Events | C (tumour TL) | C (T/N ratio) | C (host TL) |
|---|---|---|---|---|---|
| COAD | 51 | 16 | 0.677 | 0.504 | 0.658 |
| LUAD | 137 | 67 | 0.572 | 0.559 | 0.536 |
| BRCA | 115 | 18 | 0.508 | 0.521 | 0.462 |
| HNSC | 139 | 59 | 0.495 | 0.545 | 0.428 |
| STAD | 110 | 29 | 0.490 | 0.481 | 0.509 |
| UVM | 50 | 19 | 0.486 | 0.573 | 0.401 |
| BLCA | 113 | 60 | 0.485 | 0.494 | 0.458 |
| ESCA | 62 | 21 | 0.482 | 0.495 | 0.467 |
| SKCM | 131 | 109 | 0.475 | 0.487 | 0.469 |
| GBM | 50 | 41 | 0.466 | 0.455 | 0.498 |
| UCEC | 146 | 34 | 0.453 | 0.456 | 0.515 |
| LIHC | 53 | 24 | 0.452 | 0.463 | 0.459 |
| LGG | 89 | 51 | 0.434 | 0.465 | 0.413 |
| **Mean** | | | **0.498** | **0.500** | **0.482** |

Concordance was computed per cancer type in cohorts with at least 50 patients and 15 events. Chance discrimination is 0.500.

#### Table S15. Instability of CCLE lineage medians

| Lineage (TCGA code) | Cell lines | Median telomere content | 95% CI | CI width | % of spread |
|---|---|---|---|---|---|
| LIHC | 10 | 2.32 | 1.93 to 11.81 | 9.88 | 698% |
| SARC | 7 | 2.68 | 1.82 to 5.65 | 3.83 | 270% |
| ESCA | 10 | 2.60 | 2.24 to 5.66 | 3.42 | 242% |
| KIRC | 8 | 2.12 | 1.30 to 4.70 | 3.39 | 240% |
| BLCA | 7 | 3.17 | 2.01 to 5.28 | 3.27 | 231% |
| LUSC | 15 | 2.85 | 1.67 to 4.48 | 2.81 | 199% |
| LAML | 5 | 2.50 | 1.42 to 4.01 | 2.58 | 183% |
| PAAD | 11 | 2.69 | 1.27 to 3.81 | 2.55 | 180% |
| LGG | 5 | 2.06 | 1.55 to 3.75 | 2.21 | 156% |
| HNSC | 6 | 1.98 | 1.14 to 2.92 | 1.78 | 126% |
| OV | 22 | 3.39 | 2.43 to 4.02 | 1.59 | 113% |
| GBM | 8 | 2.82 | 1.66 to 3.23 | 1.57 | 111% |
| STAD | 21 | 2.33 | 1.91 to 3.45 | 1.54 | 109% |
| SKCM | 19 | 3.03 | 2.37 to 3.91 | 1.54 | 109% |
| COAD/READ | 23 | 3.09 | 1.76 to 3.29 | 1.53 | 108% |
| LUAD | 29 | 2.35 | 2.06 to 3.30 | 1.25 | 88% |
| UCEC | 11 | 2.80 | 2.42 to 3.54 | 1.12 | 79% |
| SCLC | 48 | 3.05 | 2.50 to 3.56 | 1.05 | 75% |
| BRCA | 35 | 2.18 | 1.84 to 2.58 | 0.73 | 52% |

Between-lineage spread of the CCLE medians is 1.41. A bootstrap interval wider than that carries no ranking information.

#### Table S16. Decomposition refitted by matched-normal type

| Matched normal | n | Cohorts | $\beta_{within}$ (95% CI) | $\beta_{between}$ (95% CI) | P ($\beta_b$ = 1) | P ($\beta_w = \beta_b$) | Cohort-level slope (unweighted) | SD of cohort-mean normal | Between-cohort r |
|---|---|---|---|---|---|---|---|---|---|
| Solid-tissue normal (organ-matched) | 300 | 12 | 0.420 (0.220 to 0.620) | 0.442 (0.242 to 0.641) | $7.9 \times 10^{-8}$ | 0.883 | 0.483 | 0.291 | 0.65 |
| Blood normal (reference) | 1,419 | 20 | 0.454 (0.373 to 0.535) | 1.219 (1.035 to 1.404) | 0.020 | $1.7 \times 10^{-13}$ | 1.011 | 0.164 | 0.58 |
| Blood normal (same cohorts) | 835 | 10 | 0.499 (0.407 to 0.590) | 1.272 (1.005 to 1.539) | 0.046 | $9.6 \times 10^{-8}$ | 1.103 | 0.133 | 0.75 |

Solid versus blood-on-the-same-cohorts: difference −0.831, $z = -4.90$, $P = 9.7 \times 10^{-7}$. The ratio of size-weighted between-cohort coefficients across those two arms is 2.88; for the unweighted cohort-level slope, where slope = r × SD(tumour) / SD(normal) holds exactly, it is 2.28. The predictor-spread ratio is 2.19.

The between-cohort coefficient is not a scale-free efficiency of transmission: an ordinary-least-squares slope varies inversely with the spread of its predictor, and cohort-mean blood telomere length varies far less between cancer cohorts than cohort-mean tissue telomere length does. The between-cohort *correlation*, which is scale-free, is stable across all three arms. The individual-level coefficient is likewise stable, and it is the quantity the main claim rests on.

##### Table S17. Genetic ancestry as a germline proxy, and the power available

| Term | β | 95% CI | P |
|---|---|---|---|
| Ancestry to own normal TL | 0.1403 | 0.0599 to 0.2208 | $6.4 \times 10^{-4}$ |
| Ancestry to tumour TL | 0.0384 | −0.0819 to 0.1587 | 0.531 |
| Predicted under the constitutional model | 0.0656 | — | — |

First-stage F = 11.7; instrumental-variable estimate 0.274 (95% CI −0.597 to 1.145) against an ordinary-least-squares estimate of 0.468. Power to detect the predicted effect: **19%**; patients required for 80% power ≈ 9,016, available 1,314 (117 with African ancestry above 50%).

Ancestry is not a polygenic score for telomere length. It is a germline variable correlated with the true genetic determinants and with much else besides. The validity row establishes that it carries constitutional telomere signal in this resource; the tumour-side row is uninformative rather than negative, because the predicted effect is smaller than the standard error available.

##### Table S18. The association with technical structure entered as fixed effects

| Adjustment | n | β (normal → tumour) | 95% CI | P | Partial $R^2$ |
|---|---|---|---|---|---|
| no adjustment | 1,935 | 0.532 | 0.469 to 0.594 | $1.1 \times 10^{-58}$ | 0.1263 |
| + cancer type | 1,935 | 0.417 | 0.352 to 0.483 | $1.5 \times 10^{-34}$ | 0.0757 |
| + cancer type + centre | 1,935 | 0.407 | 0.339 to 0.474 | $4.8 \times 10^{-31}$ | 0.0680 |
| + cancer type + centre + library | 1,935 | 0.407 | 0.339 to 0.474 | $4.8 \times 10^{-31}$ | 0.0682 |
| complete technical stratum (the permuted unit) | 1,935 | 0.385 | 0.316 to 0.455 | $7.6 \times 10^{-27}$ | 0.0590 |
| residual-residual Spearman (stratum-adjusted) | 1,935 | 0.280 | — | $3.1 \times 10^{-36}$ | — |

The permutation test in the main text shows that the observed pairing exceeds what shared technical structure can generate. This table gives the complementary quantity: the magnitude that survives removing that structure directly. The final model carries one fixed effect for the complete cancer-type × sequencing-centre × library-type stratum, which is the exact unit the permutation shuffled within. The last row is a Spearman correlation of the two residual series rather than a regression coefficient.

##### Table S19. Nested demographic and compositional sensitivity, including sex

| Model | n | β (normal → tumour) | 95% CI | P |
|---|---|---|---|---|
| base: normal TL + cancer type | 1,710 | 0.424 | 0.354 to 0.494 | $2.6 \times 10^{-31}$ |

| Model | n | β (normal → tumour) | 95% CI | P |
|---|---|---|---|---|
| + age | 1,710 | 0.407 | 0.335 to 0.479 | $2.4 \times 10^{-27}$ |
| + age + sex | 1,710 | 0.406 | 0.334 to 0.479 | $3.8 \times 10^{-27}$ |
| + purity + leukocyte fraction | 1,710 | 0.429 | 0.356 to 0.501 | $6.0 \times 10^{-30}$ |
| + ploidy + coverage | 1,710 | 0.421 | 0.347 to 0.494 | $2.7 \times 10^{-28}$ |
| + continental ancestry | 1,710 | 0.422 | 0.348 to 0.495 | $2.7 \times 10^{-28}$ |

Range across the ladder: 0.406 to 0.429, a spread of 0.023. Sex is itself a determinant of the matched normal in this set (male versus female, β = −0.047, P = 0.013), so the stability above is not an absence of a sex effect.

#### Supplementary Notes

##### Note 1. Barcode harmonisation and the merge procedure

TCGA identifiers appear at four levels of resolution in the source files used here: 12-character patient barcodes, 15-character sample barcodes, 16-character sample-plus-vial barcodes and 28-character aliquot barcodes. The telomere table is keyed at aliquot level, ABSOLUTE purity at sample level, and the immune-landscape and clinical releases at patient level. Merging without harmonisation silently drops the majority of rows.

All identifiers were therefore truncated to the level of the coarsest key required by a given join, and sample-type codes (TP, TM, TB, NB, NT) were parsed from positions 14–15 of the barcode rather than taken from any free-text annotation. Joins were performed at sample level where both sides carry a sample identifier and at patient level otherwise, and every join was audited for the number of rows entering and leaving.

Match rates are reported in Supplementary Fig. S1d. Primary tumours match at 93–97% to all three external annotations. Normal samples match at 0%, which is correct and expected: ABSOLUTE purity, leukocyte fraction and CIBERSORT fractions are defined for tumour samples only. This was verified rather than assumed, because a 0% match rate is otherwise indistinguishable from a broken merge.

##### Note 2. Why the analysis set is restricted to whole-genome and low-pass data

The restriction is empirical. Telomere attrition with age in leukocytes is one of the most robustly replicated quantitative facts in telomere biology, and any estimator that measures telomere length in blood should recover it. Whole-genome and low-pass estimates do (Supplementary Tables S7 and S7b). Whole-exome estimates fail to recover it: the unadjusted slope is positive in blood normals, in solid-tissue normals and in primary tumours alike, and once cancer type, sequencing centre and sex are adjusted for it is indistinguishable from zero rather than reversed.

Two candidate explanations were considered. The failure was not reproduced in the 36-line CCLE comparison (main text Fig. 2f), which shows that the discordance is not universal to exome sequencing but does not identify what makes clinical material different: admixture, specimen or DNA quality, technical batch, capture protocol and their combinations are all consistent with the observation and are not separated here. Classical random measurement error would attenuate a slope toward zero rather than reverse it, which is what the adjusted and within-cohort exome estimates in fact show; a systematic, composition-dependent bias could produce either, so the unadjusted positive sign should not be over-read. Because exome data are 78.6% of the resource, restricting to whole-genome and low-pass libraries removes most of the samples — but retaining them would mean averaging a calibrated measurement with one that is demonstrably not measuring the intended quantity.

An important consequence: the cross-cancer ranking derived from exome data does not agree with the ranking derived from whole-genome and low-pass data. Any re-analysis of this resource that pools assays inherits that disagreement.

**Note 3. Circularity among tumour-purity estimators**

Tumour purity has no single ground-truth measurement, and the available estimators derive from different data modalities: copy number (ABSOLUTE) [38], bulk expression (ESTIMATE) [33], leukocyte-specific unmethylation (LUMP), histological review (immunohistochemistry), and a consensus of the four (CPE) [32].

This matters here because the immune covariate used throughout — leukocyte fraction — is itself methylation-derived [34]. LUMP and leukocyte fraction are estimated from overlapping methylation signal, and CPE contains LUMP. Regressing telomere length on CPE while adjusting for leukocyte fraction therefore places two partially identical quantities in the same model, and the resulting coefficient is not interpretable as an independent purity effect.

Supplementary Table S11 quantifies this. The correlation between each estimator and leukocyte fraction ranges from −0.87 (LUMP) to −0.28 (immunohistochemistry), and the apparent purity effect on telomere length tracks that ordering almost exactly, roughly doubling from ABSOLUTE to ESTIMATE. ABSOLUTE was used as the primary purity variable for this reason. The substantive conclusions do not depend on the choice — the cross-cancer ranking and the host effect are robust under either — but the magnitude of the reported purity effect does, and papers that report a purity effect using a consensus or methylation-derived estimator alongside methylation-derived immune covariates should be read with this in mind.

**Note 4. Verification procedures: figure auditing and citation checking**

Two automated checks were built because manual inspection had already failed at least once in each domain.

**Figure auditing.** Each figure script is re-executed with the export function intercepted, and the live figure object is inspected before rendering. The auditor tests, for every panel: pairwise bounding-box overlap of all text artists within a panel and across panels; text extending beyond the figure canvas; tick labels rendered as phantom values ("nan", "None", "<NA>"); and axes that mix incompatible units on one scale. Only tick labels inside the current view limits are considered, since matplotlib retains formatter output for ticks it does not draw. The auditor reports 0 issues across all 10 figures. It was written after a reader identified two text collisions in Supplementary Fig. S1 that visual inspection had missed; re-running it then found the same defect class in four further figures.

**Citation verification.** Every reference is resolved by explicit identifier — PubMed identifier or DOI — rather than by title search, and the retrieved title is checked against the expected title before the record is accepted. The identity check runs first, prior to any cross-source comparison, because agreement between two databases about the wrong paper is not evidence of anything. Records whose titles begin with a correction, erratum, comment or peer-review prefix are rejected even when the remaining title matches exactly, since correction notices and review reports reproduce the original title verbatim. Reverse token coverage is required so that a document whose title merely contains the expected one is rejected, while an appended subtitle is permitted. Each accepted record is then confirmed independently against Crossref. Of the 57 references, 56 were confirmed by both sources and 1 by PubMed alone, the remainder having no Crossref record that matched on identity.

The manuscript compiler refuses to emit output if any citation key in the text is absent from the verified set, or if any citation marker survives substitution. An unverified reference cannot reach the compiled document.

**Numeric claims.** A screen that asks only whether a number appears somewhere in the results directory was measured against random input and found to pass essentially every three-decimal value between 0 and 1; it therefore carries no information for correlations and coefficients, and is retained only as a coarse filter that reports its own false-pass rate. The load-bearing check is a pinned manifest in which each of 94 headline claims names the exact source file and cell it derives from, the precision at which it is reported, and the exact string that must appear in the text. A claim fails if the source no longer yields that value or if the string is no longer present, so the check catches both an analysis that has changed under the prose and prose that has drifted from the analysis. All 94 verify. Numbers quoted from other publications cannot be checked this way and are instead recorded, with verbatim quotations and their sources, in a separate external-claims file.
